# The Once and Future Fish: 1300 years of Atlantic herring population structure and demography revealed through ancient DNA and mixed-stock analysis

**DOI:** 10.1101/2024.07.11.603078

**Authors:** Lane M. Atmore, Inge van der Jagt, Aurélie Boilard, Simone Häberle, Rachel Blevis, Katrien Dierickx, Liz M. Quinlan, David C. Orton, Anne Karin Hufthammer, James H. Barrett, Bastiaan Star

## Abstract

Atlantic herring populations have been the target of highly profitable coastal and pelagic fisheries in northern Europe for well over a thousand years. Their complex and intermingled population dynamics have sparked extensive debate over the impacts of historical overfishing and have complicated their sustainable management today. Recently developed tools – including diagnostic SNP panels for mixed-stock analysis – aim to improve population assignment for fisheries management, however, the biological relevance of such tools over long periods of time remains unknown. Here, we demonstrate the millennium-long applicability of diagnostic SNP panels and identify population perturbations associated with increasing exploitation pressure and climate change by analyzing whole genome data from modern and ancient herring specimens. We find that herring demographic cycles were likely within healthy ecosystem boundaries until the dramatic disruption of these cycles in the 20th century. We find only autumn-spawning herring in our archaeological remains spanning 900 years from 8 sites across Europe, supporting observations that the numerical dominance of specific spawning populations can demographically outcompete other herring types. We also obtain pre-archival aDNA evidence for the famous, cyclical “Bohuslän periods,” during which mass quantities of North Sea autumn-spawning herring congregated in the Skagerrak. Finally, the long-term applicability of diagnostic SNP panels underscores their highly cost-effective application for the genetic monitoring of herring stocks. Our results highlight the utility of ancient DNA and genomic analysis to obtain historical and natural insights in herring ecology and population dynamics with relevance for sustainable fisheries management.

## Introduction

### Herring Fisheries Past and Present

Atlantic herring (*Clupea harengus*) have been a crucial source of food and economic activity in the nations surrounding the North Sea for well over a thousand years (1–3). Historical fishing operations in the North Sea likely targeted the North Sea autumn-spawning herring, which, until the 1970s, supported one of the largest herring fisheries in the world (4,5). In England, fishers would migrate down the coast of the North Sea following spawning aggregations; herring was so abundant c. 1000 CE that it was called “herringsilver” and used as payment in tithes and taxes (2). The marked increase in herring remains in archaeological sites in England following 1000 CE, alongside the appearance of large quantities of other marine fish species, has led archaeologists to suggest — somewhat tongue-in-cheek — that this date represents the “Fish Event Horizon” (FEH); a point at which marine populations were exploited at scale for the first time in this region (6). North Sea herring were also the source of the famous “Bohuslän Periods,” a phenomenon lasting from the 10th-19th centuries, in which enormous schools, influenced by climatic cycles, would suddenly migrate to new spawning grounds in the Skagerrak (7–9). During these 20-50 year cycles, entire herring industries would spring up around the spawning migrations only to disappear once the herring shifted back to the North Sea (8,9). Perhaps the most famous — and most extensively studied — herring fishery in the North Sea was that of the Dutch Republic. Having developed technology that allowed for an unprecedented high seas fishery, the Dutch Republic dominated European herring trade in the 16th-18th centuries, peaking between the late 1500s and 1600s (4,10). Fishing expansion in Northern Europe during this time was so extensive that it represents a second phase of commercialization (11). Herring was then a profitable commodity and the Dutch Republic controlled up to 80% of the market, resulting in an influx of wealth that supported the “Dutch Golden Age’’ of scientific and political experimentation as well as global exploration and colonization (4). Herring was so important to the Netherlands it was called “Gratia Dei” (“gift from god”), a “golden mountain,” and the “triumph of Holland” (12).

By the 20th century, herring stocks throughout the eastern Atlantic were heavily exploited with little regulation, resulting in mass stock collapses across the species range (13). Despite warnings from the International Council for the Exploration of the Sea (ICES) Herring Assessment Working Group (HAWG), management bodies failed to reduce Total Allowable Catch (TAC) in the region until the population had been nearly entirely wiped out (5). By this time, North Sea Autumn Spawners spawning-stock biomass (SSB) is estimated to have declined by 97%, from 5 million tonnes in 1940 to 50,000 tonnes in 1970 (5). By 2010, SSB had recovered to 1.5 million tonnes (5,14), but this success was shortly followed by recruitment failures again in recent years (15). The North Sea herring fishery currently accounts for 21% of the total economic value of the EU pelagic fisheries sector, thus ongoing recruitment failures will have widespread impacts on the EU economy as well as the North Sea ecosystem (16). In recent years, ICES has again recommended reductions in TAC for the North Sea Autumn Spawners (NSAS), as recruitment continues to falter (15). Landings for fisheries in the North, Celtic, and Irish Seas (the regions encompassed by sampling in this study) are consistently higher than the recommended TAC, although stocks are assessed as in varying degrees of health (15).

A key aim in determining sustainable management policies is establishing what constitutes healthy ecosystem thresholds (17). Understanding ecosystem thresholds necessitates acknowledging that the region has been exploited for over 1000 years, thus efforts must take historical exploitation levels into account rather than relying on landing records that begin in the 20th century — a time at which marine resources were likely already heavily depleted (18). Historians modeling catch records initially concluded that historical herring industries could not have impacted herring stock size (4), although recent research suggests landings across Europe were far larger than previously understood (10). Assessing the impact of historical fisheries, historical trade, and shifting fishing pressures hinges on correctly identifying the biological populations targeted in the past. While the dominance of North Sea fisheries after the 16th century in the European commodity market is well characterized, we know little of the relative contribution of the North Sea fisheries to the European commodity herring trade around 1000 CE.

Historical records show that herring were traded from England to the European continent around this time (11), nevertheless the extent of this trade is unknown. Questions remain regarding the quantity and range of the English herring trade, with many herring remains from archaeological sites around Europe that pre-date the peak of the Dutch herring period assumed to stem from the Øresund fishery, which dominated the market from 800-1200 CE (19–21). Determining the relative contribution of these industries is essential for our understanding of European history as well as correctly modeling our historical impacts on the ocean. The application of ancient DNA provides novel precision in determining population of origin for archaeological remains (22–24). Identifying population of origin will allow for direct comparison of samples across time, assessment of population continuity, and —by including historical and archaeological context— identification of previously-unknown routes of trade (25–27).

### Complex population dynamics of Atlantic herring

Biologically distinct herring stocks mix during migration (e.g., during feeding phases) and modern fishing practices therefore often result in mixed-stock exploitation (28). Mixed-stock exploitation has increased over time as technological advances allowed fishing operations to target increasingly distant fish schools, such as feeding aggregations, as opposed to fisheries in the Middle Ages that were often restricted to targeting biological stocks via coastal spawning aggregations (2,21). Exploitation of mixed stocks has complicated efforts to introduce sustainable fishing practices (29); expert policy advice recommends management based on biological populations rather than a mixed stock to improve sustainable exploitation (15). Importantly, marine fish populations do exhibit biologically meaningful fine-scale population structure, despite strikingly low intraspecific F_ST_ values (30). Thus, novel genetic techniques have been developed for mixed-stock analysis (MSA) of Atlantic herring with the aim that these can at some point be applied to a revision of current ICES subdivisions and recommendations, and perhaps provide fishers with onboard tools for real-time management decisions (28,31). These tools (panels of single-nucleotide polymorphisms (SNPs) known to be different between populations) have been demonstrated as highly capable of assigning an individual to the biological populations studied, even in those cases with low overall population differentiation. For instance, based on such tools the English Channel (Downs) and North Sea Autumn Spawning (NSAS) populations were divided into separate ICES management units in 2022 (15,31–33). However, many of these tools are still under development and it remains unknown how well these tools perform over long periods of time or if diagnostic SNP panels used for MSA will need to be reassessed frequently (28).

In recent years, the application of assignment tools to ancient specimens has demonstrated population structure continuity on the level of metapopulation for several marine fish, including Atlantic cod (34) and Atlantic herring (20). Previous ancient DNA studies for Atlantic herring have focused on assigning population to relatively broad metapopulations based on spawning season and local adaptation (e.g., salinity levels) (20). Mixed-stock analysis techniques – which use diagnostic SNP panels to identify the biological population of fish that are exploited together – have thus far not been applied to ancient fish remains on a pan-European scale. Understanding how well these diagnostic SNP panels work over time will facilitate their development and implementation by providing information on their cost-effectiveness and the underlying evolutionary processes that provide their statistical power and utility. For example, the validity of diagnostic SNP panels for deep time series data implies that these sites in question are under strong pressure to maintain population structure and local adaptation. Such findings would show that the creation of diagnostic SNP panels is highly cost effective as a management strategy — as they could be used for a long time — and would highlight the need to monitor these populations as genetically unique, providing the rationale for individual management units. In contrast, deviation from such population continuity would thus also suggest strong perturbation on these herring stocks, indicating cause for stricter management and/or consideration of changing environmental impacts.

We here use ancient DNA and whole genome analysis to assess spatiotemporal changes in stock exploitation of herring in the North Sea, English Channel, and Celtic and Irish Seas. Archaeological herring specimens spanning the 8th-16th centuries were collected from sites in England, Norway, Switzerland, and the Netherlands. We demonstrate changes in herring genetic diversity and population structure over time and assess anthropogenic and climatic forcing on herring demographic trends. We discover that modern diagnostic SNP panels designed for mixed-stock analysis in sustainable management regimes have deep-time relevance by applying mixed-stock analysis to archaeological herring remains. Finally, we recover historically-reported phenomena — such as the known stock collapses in the 20th century and the cyclical “Bohuslän periods” — by comparing ancient and modern whole-genome data.

## Results

### Population Structure & Mixed-Stock Analysis (Contemporary)

Atlantic herring are structured into metapopulations by adaptation to different salinity levels and spawning season based on PCA analysis of high-coverage whole genome data of 68 specimens from across Europe (Figure 1) (20,35). Atlantic and Baltic herring cluster separately with autumn/winter spawners segregating from spring spawners within these two clusters (Figure 1a). An additional cluster lying outside of this grouping comprises Transition Zone populations, which spawn in the spring and live across the salinity gradient between the North and Baltic seas. The herring stocks under consideration in this study – British, Irish, and North Sea Autumn/Winter spawners (BINSA) – clustered together in one metapopulation on the PCA (Figure 1b). This metapopulation can be split into three subpopulations: Isle of Man, Celtic/Downs, and North Sea (NSAS) based on mixed-stock analysis (MSA) using 82 SNPs that discriminate between herring populations in the North Atlantic (28,31). MSA suggests the North Sea subpopulation is more distinct from the other three subpopulations (Celtic Sea, Downs, and Isle Of Man), and that Downs and Celtic Sea populations are unable to be distinguished based on this SNP panel (Figure 1c,d), which is in line with previous studies (31).

**Figure 1.**
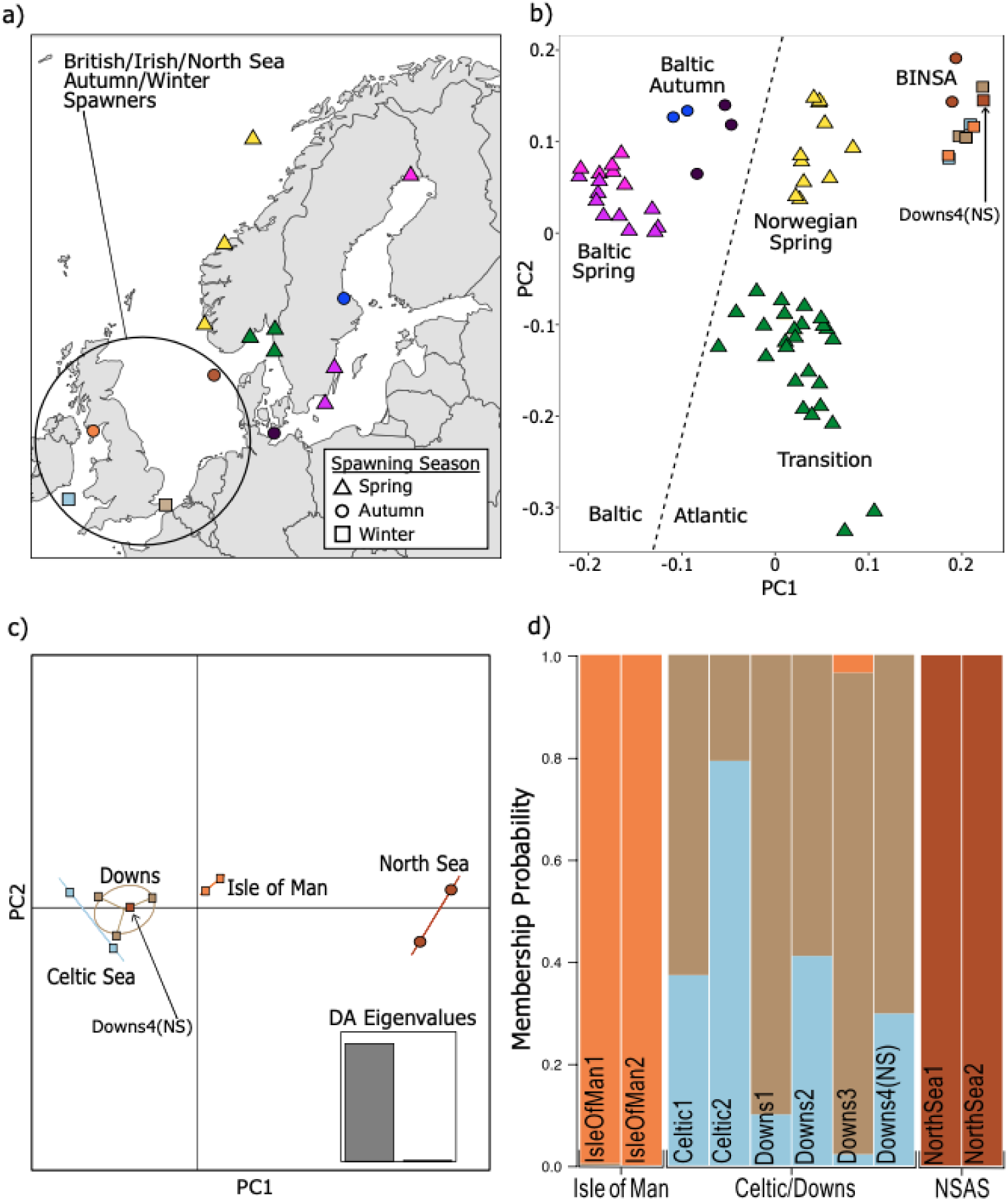
Population structure in modern Atlantic herring is based on local adaptation. a) Sampling locations for modern herring samples. Shapes correspond to spawning season; b) Population structure is driven by adaptation to salinity (Baltic vs Atlantic) and spawning season. Samples are colored by sampling location (panel a) and shapes correspond to spawning season. PCA constructed with 10 million SNPs (non-maf filtered) using smartPCA; c) Mixed-stock analysis with the BINSA metapopulation shows substructuring between the North Sea, Isle of Man, and Celtic/Downs populations, as previously reported (28,31). This panel shows the results of DAPC analysis using two PCs; d) Three subpopulations are indicated by DAPC analysis: North Sea, Isle of Man, and an admixed population between Celtic Sea and Downs based on 82 SNPs diagnostic SNPs designated for mixed-stock analysis (28,31). Four populations were used as input priors: Celtic Sea, Downs, Isle of Man, and North Sea (NSAS). DAPC suggests there are three groups: Downs4(NS) was identified as an admixed Celtic Sea/Downs individual, not a North Sea individual as listed in public metadata. Samples are grouped based on genetic identity (labeled on x-axis). Labels within bars refer to individual sample IDs.

One of the modern individuals listed as sampled within the NSAS population showed high similarity to the Downs/Celtic Sea population (Figure 1b; Figure S1). This sample was not collected in a region where NSAS typically spawn, but in a region associated with the feeding migration (35,36). We therefore considered this a mixed-stock sample and re-classified this individual as genetically belonging to the “Downs” population in all subsequent analyses (Downs4(NS)). Substructure analysis based on discriminant analysis of principal components (DAPC) (37) shows that the optimum number of PCs to describe the modern data is 1 (Figure S2) and the DAPC find.clusters function to determine the number of true populations suggests K=2, indicating the most significant population structure is exhibited between the NSAS and all other BINSA subpopulations. Winter spawners south and west of the UK have a monophyletic relationship based on Treemix analysis of the diagnostic SNP panel (Figure S3). A single migration event from the common ancestor of the western BINSA populations to the North Sea was detected using OptM (38) (Figure S3, S4). Bootstrapping revealed lack of consensus on the placement of the edge, thus the consensus tree shows gene flow from the MRCA of Isle Of Man, Downs, and Celtic Sea to the contemporary North Sea population. No population differentiation between BINSA subpopulations was found using Treemix (40) based on whole genome data. This result corroborates that herring population structure is restricted to a few highly-differentiated genomic regions (Figure S5).

### Archaeological Specimen Assignment

32 archaeological specimens from 8 different sites dated to the 8th-16th centuries were assigned to metapopulation by determining their adaptation to differing salinity levels (e.g., Baltic vs Atlantic) and spawning season with BAMscorer v1.5 (24,39). All ancient specimens were autumn spawners; no spring spawning herring were identified in the archaeological sites (Figure 2). Archaeological remains from Basel, Switzerland, (Schnabelgasse 6 and Museum der Kulturen, Figure 3) (40), showed mixed origins, with some samples stemming from the Baltic and some from the Atlantic. All other sites exclusively contained samples that were assigned to the modern-day BINSA metapopulation. Eight archaeological specimens had sufficiently high coverage to conduct DAPC analysis and assess the ability of MSA SNP panels to be used over long timescales. These samples stemmed from the sites of Lyminge in southeastern England (8th-9th century CE) (41), Coppergate (930-1000 CE) (42) and Blue Bridge Lane (13th century) (43) in York, and Huis in den Struys (15th century) (44) in the Netherlands. All high coverage samples except HuisDenStruys2 were assigned to the winter-spawning population from the southern and western sides of the UK (Figure 3a). A higher degree of similarity between Isle of Man and the Celtic Sea/Downs population is exhibited in the ancient samples than the modern samples (Figure 4). The classification based on diagnostic SNPs (28,31) is consistent across the entire time period studied, with the largest genetic divergence consistently identified between the North Sea population and the other three subpopulations (Jost’s D (45), Figure 3b). These results agree with DAPC analysis on the contemporary samples showing two main subpopulations within BINSA: North Sea autumn spawners and southern and western winter spawners with some degree of substructure between Celtic/Downs and Isle of Man. Genetic divergence measured in modern populations alone suggests additional structure between Celtic/Downs and Isle of Man (Figure S6).

**Figure 2.**
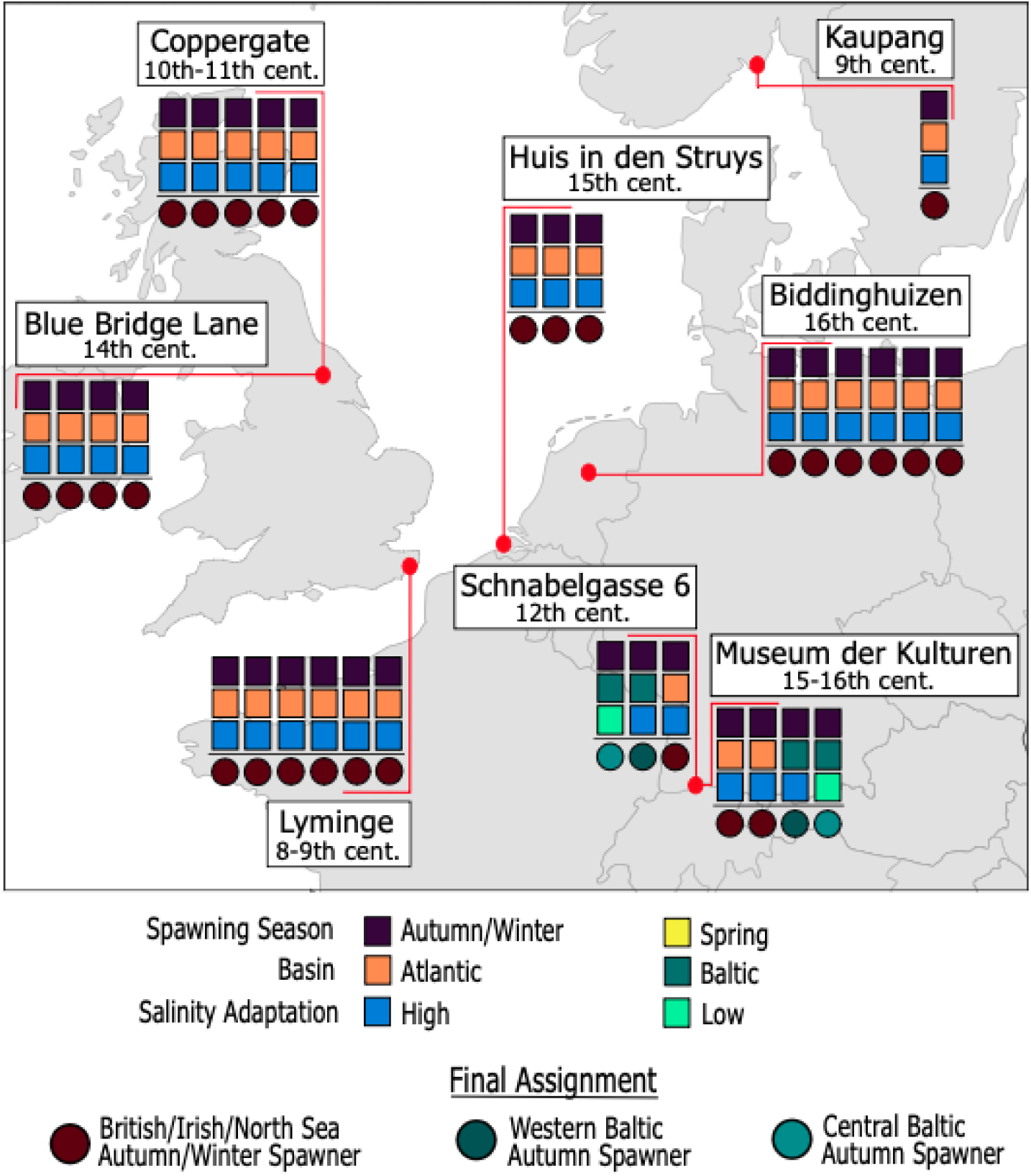
Metapopulation assignment for archaeological remains shows all archaeological herring are autumn spawners. Two samples each from Schnabelgasse 6 and Museum der Kulturen stem from the Baltic, whereas all other samples are from the BINSA metapopulation. Biological population assignment using BAMscorer for low-and high-coverage ancient herring remains. Samples are assigned to a final metapopulation through three hierarchical assignment tests. Each square represents the results of one assignment test: spawning season (spring vs autumn/winter), ocean basin (basin can be determined from the chromosome 12 inversion for autumn spawners), and salinity adaptation (high/low/mid). Circles represent the final BAMscorer assignment according to the combined assignment tests. Each column corresponds to one specimen.

**Figure 3.**
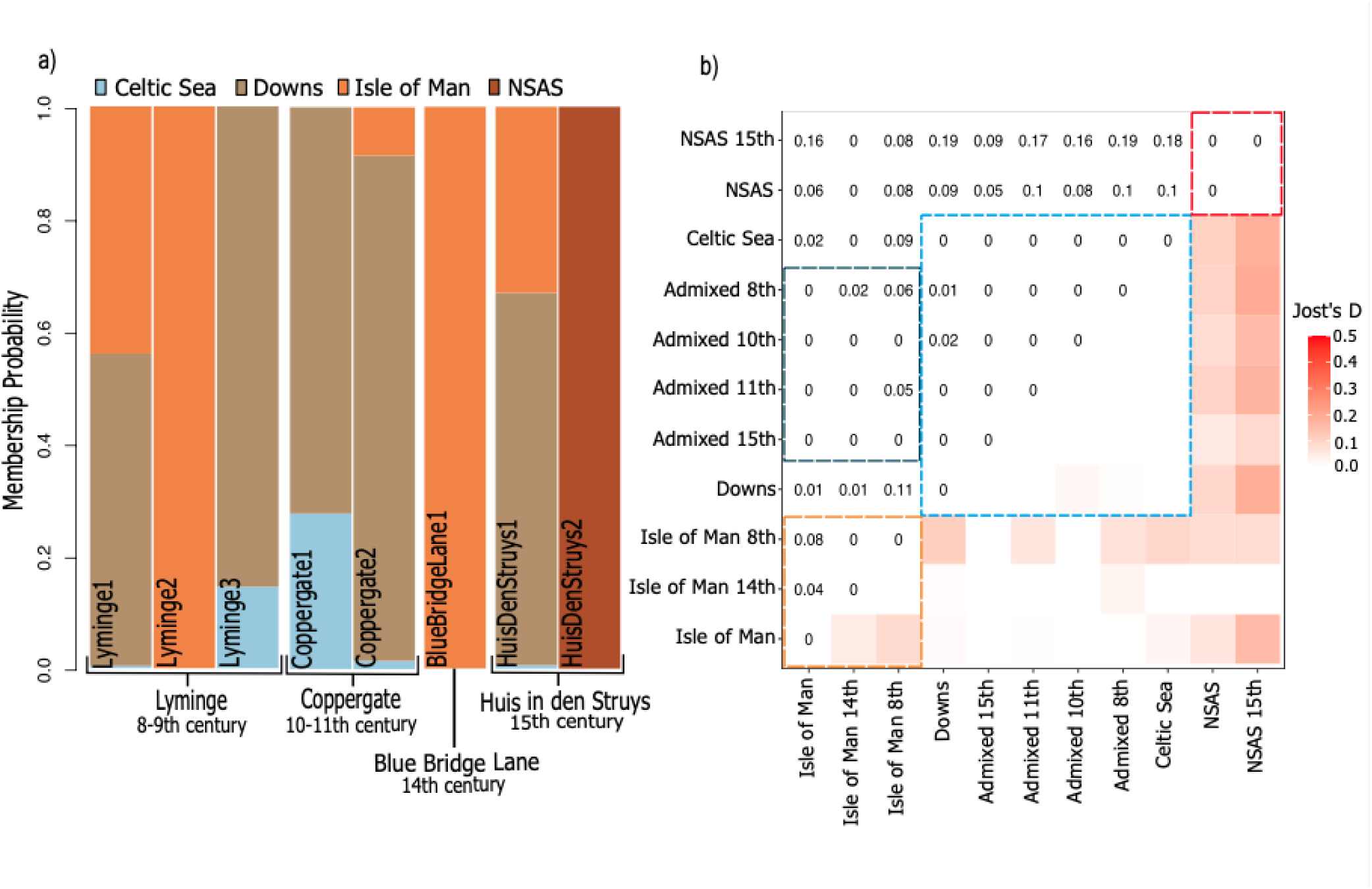
Mixed-stock analysis of high-coverage ancient specimens shows long-term population substructure in the BINSA metapopulation. a) Archaeological samples are assigned to each of the three subpopulations identified for the contemporary samples: Isle of Man, North Sea, and Celtic/Downs. The ancient remains, however, show a much greater degree of genetic admixture between Downs and Isle of Man. Colors correspond to input population priors based on sampling locations for contemporary stocks; b) Persistent population substructure across time is shown through absolute divergence analysis based on subpopulations separated by time depth. Intrapopulation Jost’s D throughout time largely shows values of 0 for Celtic/Downs and North Sea. Higher Jost’s D is exhibited within IsleOfMan over time, whereas IsleOfMan samples show more similarity with ancient Celtic/Downs populations (dark blue hashed box), likely reflecting increased admixture with Celtic/Downs in the past. Labels refer to populations separated by time period to account for possible change over time. Hashed lines show the three major populations: IsleOfMan (orange), Celtic/Downs (blue), and NorthSea (red). “Admixed” refers to the ancient individuals that showed admixture between Isle of Man, Celtic Sea, and Downs in varying degrees.

**Fig 4.**
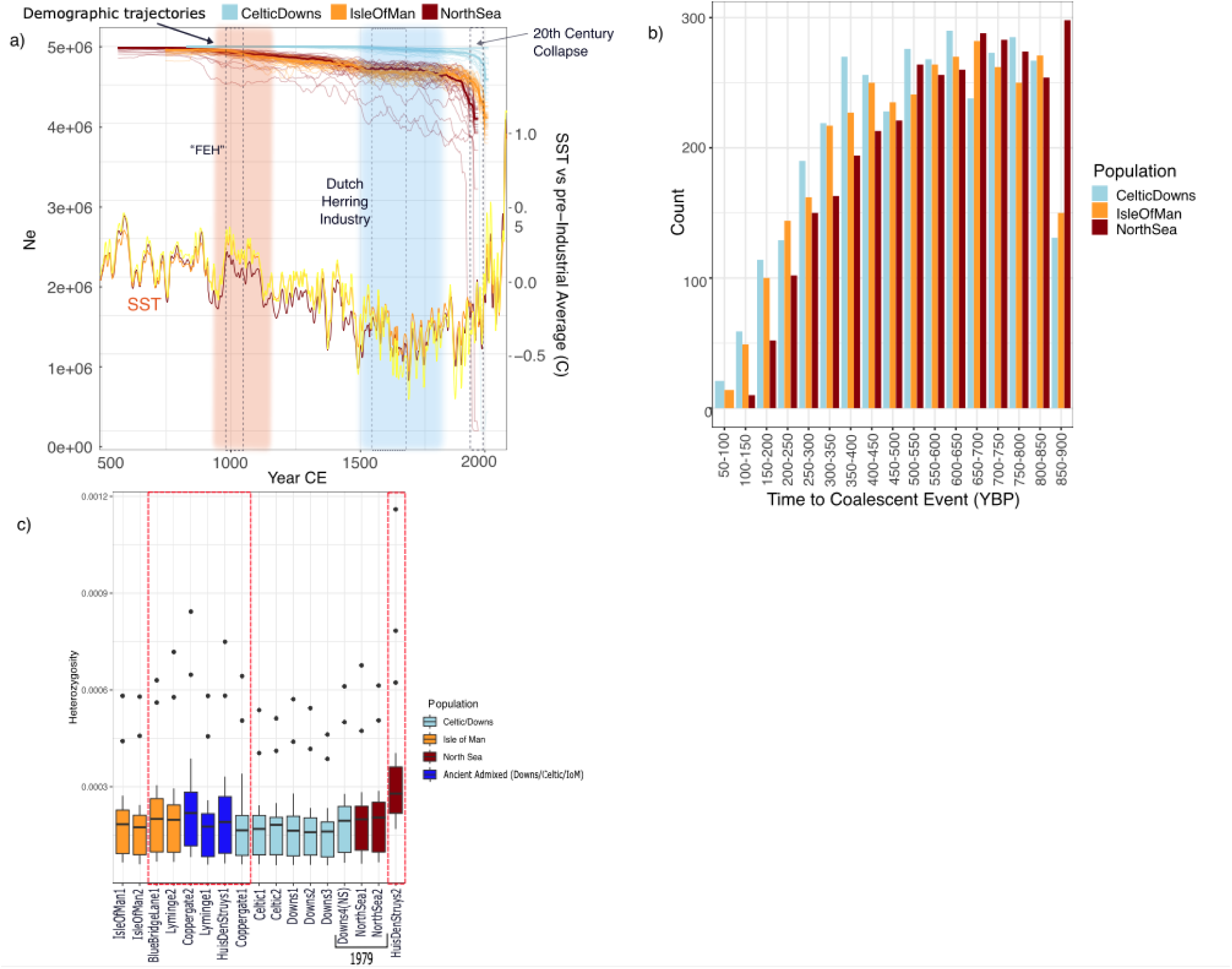
Genetic diversity and demographic change over time and space suggest recent population declines in exploited herring stocks. a) N_e_ reconstructions showing recent dramatic declines, with population declines beginning ∼1000 CE for NSAS and Isle of Man. N_e_ is estimated per population with GONE. Opaque lines represent median N_e_ estimates from 30 iterations of GONE, with each iteration represented by semi-transparent lines. Year was calculated by scaling generational N_e_ estimates to generation times reported by Feng et al. (99). Colored blocks represent the Medieval Climate Anomaly (orange) and Little Ice Age (blue). Hashed blocks represent historical events. SST reconstructions were adapted from three published indicators (68). b) Number of ROHs associated with a coalescent event at a specific time period (calculated with the formula T=100/2*cM * Gen) per modern population. c) Higher heterozygosity in ancient/historical samples (red hashed boxes) than 21st century samples . Heterozygosity was estimated per chromosome by individual using the site frequency spectrum. Samples are grouped according to site/basin of origin and boxplots are filled according to population assignment from MSA.

### Changes in Genetic Diversity Over Time

Ancient samples exhibited larger variation in heterozygosity estimates than modern samples and a slightly higher median heterozygosity estimate per individual when compared to the contemporary samples. HuisDenStruys2 (North Sea) exhibited a significantly higher heterozygosity estimate than the North Sea samples collected in 1979, which in turn exhibited higher heterozygosity than the samples collected in the 2010s (Fig 4c). Runs of homozygosity (ROH) were also assessed per population. The contemporary populations exhibited a pattern of ROH that suggests increased inbreeding throughout history, with the majority of individuals exhibiting the highest number of ROH with estimated coalescent ages between 600-850 YBP (Figure 4b; Figure S7). The North Sea population diverged from this pattern by exhibiting the largest number of ROH with coalescent ages estimated to 850-900 YBP, a time bin for which the Celtic/Downs and Isle of Man populations exhibit the smallest number of ROH, suggesting a significantly higher level of inbreeding for the NSAS at this time. The only ancient samples with high enough quality for ROH analysis were from Lyminge and had been assigned to separate populations with MSA (Lyminge2, Lyminge3, see Figure 3a), therefore they were excluded from population-wide ROH analysis. The fraction of the genome in ROH between ancient and contemporary samples was thus compared on an individual basis. Lyminge2 (Isle of Man) and Lyminge3 (Celtic/Downs) were compared with the randomly- downsampled modern individuals used for the population comparison. Both ancient samples had smaller total lengths of ROH across their entire genomes and a lower fraction of the genome in ROH than the contemporary samples (Table 1). Notably, Lyminge2 exhibited less than half the number and total length of ROH than Lyminge3.

**Table 1.**
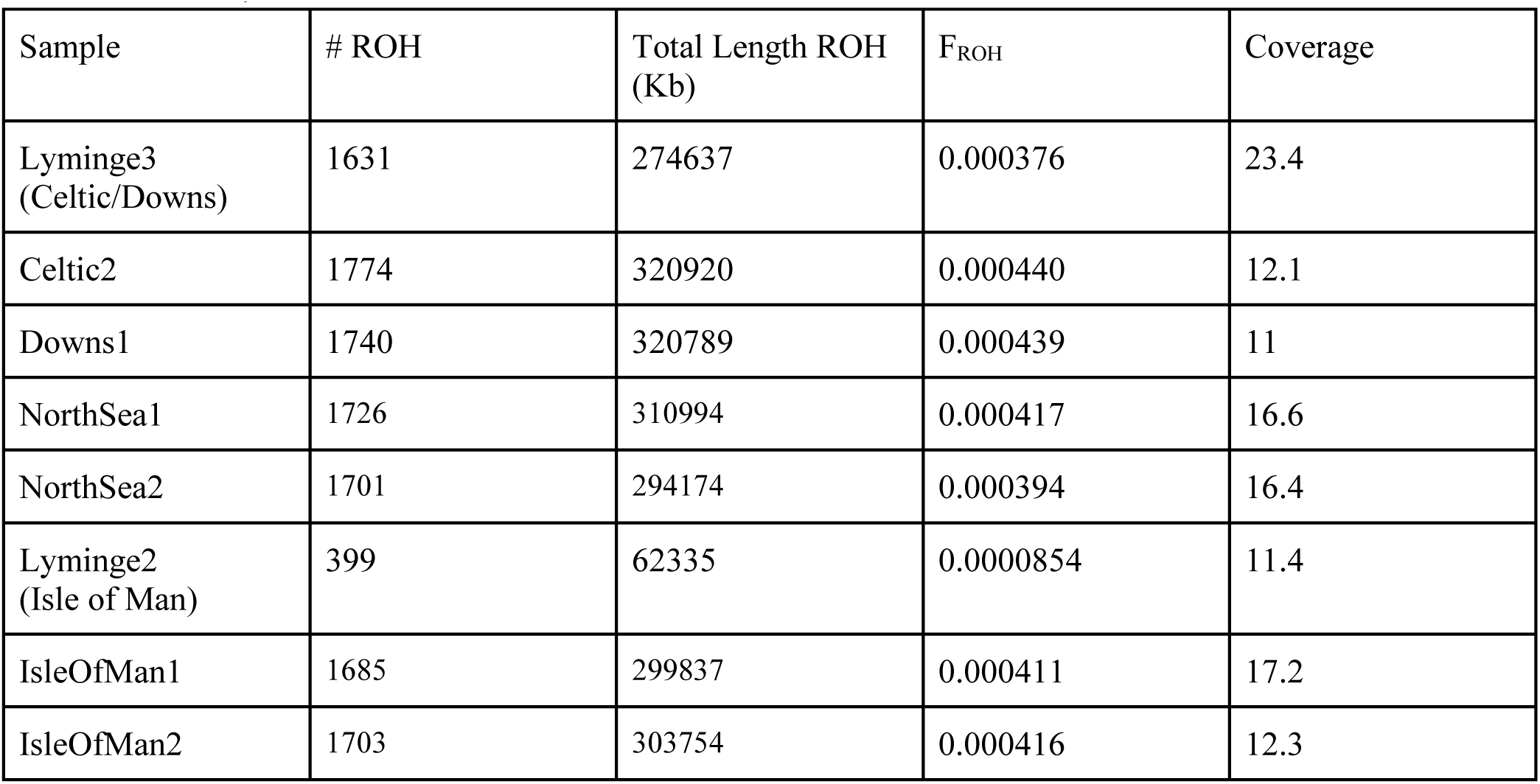
F_ROH_ by individual.

### Demographic Modeling

When all herring were grouped as a single population, demographic modeling using GONE (46) exhibits two alternating demographic patterns (see Figure S8) and characteristics that may indicate past admixture (47). GONE is known to be highly sensitive to population structure (46,48,49), indicating the population substructure discovered through MSA analysis is impacting this analysis. Given the time difference in sampling (50 years) between some of the subgroups and the demonstrated substructure within BINSA from DAPC mixed-stock analysis, all four subpopulations were run separately. When run following original metadata (e.g., North Sea, Celtic Sea, Downs, and Isle Of Man), North Sea, Downs, and Celtic Sea population showed signs of admixture (Figure S9). Re-segregating samples based on their classification with DAPC removed these admixture artifacts, further supporting population assignment results. Downs4(NS) was removed from this analysis as it was sampled 50 years prior to the remaining Celtic/Downs samples. All BINSA subpopulations showed a similar demographic trend, with the North Sea population showing a larger degree of variation in N_e_ estimates across all iterations compared to the other three subpopulations. All three subpopulations have relatively stable populations prior to the 20th century, albeit with a slow decline beginning around 1000 CE (Fig 4a). We observe consistent signatures of decline in all three subpopulations beginning in the mid-20th century, which matches both known commercial collapse (5,14) and increasing SST (Figure 4a). No population shows signatures of recovery in recent decades.

## Discussion

We have here demonstrated that the existing population substructure identified among modern herring populations in the North Sea, Channel, and Celtic and Irish seas persists across centuries. We further find that herring demographic patterns were likely within ecosystem boundaries until the dramatic 20th century disruptions. The long-term persistence of population structure in the Atlantic herring suggests mixed-stock analysis (MSA) based on SNP panels is a cost-effective technique that identifies evolutionary distinct biological populations for sustainable management. Recent perturbations in herring demography further indicate that anthropogenic impacts on the species, both in terms of exploitation and climate change, are greater than previously understood. In the following paragraphs, we discuss the novel historical and biological insights, and management implications of our results.

### Historical & biological implications of archaeological population assignments

Our archaeological assignments lead to spatiotemporally broad as well as local historical insights. On a European scale, the vast majority of modern herring landings in the North Atlantic and the Baltic stem from spring-spawning herring stocks (50,51). Historical herring exploitation in the Baltic however, relied exclusively on autumn-spawning herring until the mid-20th century (20). Here, we discover that all 32 archaeological herring specimens from 8 sites across Europe (Norway, Switzerland, the Netherlands, England), spanning a period of 900 years (700-1600 CE) consist of autumn spawning herring only. In particular, the assignment of the archaeological remains from Switzerland – which were obtained through diverse sources of long-distance trade – strengthens the argument that only autumn-spawning herring were traded as a commodity product in Europe until relatively recently. Intriguingly, it appears that Atlantic ocean basins are dominated by one spawning phenotype at a time (50); for instance, population declines in autumn-spawners resulted in dominance of spring-spawners in the Baltic (20,50). Although there are small, coastal populations of spring spawners in the North Sea (5), our results imply a long-term dominance of autumn- and winter-spawning herring over hundreds of years despite known perturbations from climate and exploitation in the Baltic, North Sea, and Irish/Celtic Sea regions.

Detailed assignment results using mixed-stock analysis from eight archaeological specimens provided further insight into past herring biology. What is striking in our results is the presence of mixed assignment between the Celtic, Downs, and Isle of Man populations in significantly greater proportions than found in contemporary samples. This pattern is reflected across all archaeological sites, from the 8th to 15th centuries. These results suggest admixture or gene flow between these subpopulations that existed in the past but are currently not detectable. Our results indicate increased gene flow between these populations as recently as the 15th century, likely mediated through the Celtic Sea population. Herring are known to exhibit the “basin effect” – retreating to core habitats during population decline and re-colonizing larger ranges only at large population sizes (5,13,52). Thus, increased admixture in the past suggests larger population sizes for the Celtic/Downs and Isle of Man populations than today, where admixture does not appear. Changed herring subpopulation connectivity has implications for historical interpretation of population assignments. Archaeological population assignments are here discussed in the context of each archaeological site.

### Switzerland

Basel comprises two archaeological urban sites: Schnabelgasse (12th century), and Museum der Kulturen (15th- 16th centuries). Basel was at the time, a distribution center for commodity goods such as herring (53). The majority of these specimens stem from the Baltic, likely traded from the Øresund industry and its successors (20). There are various trade routes these specimens could have taken, including being imported to England and then re-exported to the continent, or through trade routes along the Rhine (53). Genome sequence quality of samples assigned to BINSA was not high enough to identify their subpopulation. Given the known long-distance trade from southern England to France and Germany (11) it’s possible the BINSA remains stem from fisheries along the south or east coast of England. This origin is particularly likely for the remains stemming from the 12th century, at which time herring was traded from southern England to the continent for wine from France (1), and as the Dutch herring industry was not yet dominant (4). The samples recovered in Basel from the 15th-16th century however, likely do stem from the North Sea autumn-spawners and the Dutch herring industry. Overall, the herring remains from Basel highlight the overwhelming presence of autumn-spawning herring in the commodity fish trade during the medieval era.

### Norway

Kaupang is one of the earliest Viking urban centers in Scandinavia, dating to the 9th century. It is located in southern Norway on the Skagerrak (54). Today, herring in the Skagerrak consist almost entirely of Norwegian spring-spawners and western Baltic spring-spawners; while NSAS are today sometimes found in the Skagerrak, they do not appear in large, coastal aggregations (55). Thus, the identification of the Kaupang herring sample as an autumn- rather than a spring-spawner is unexpected. NSAS coastal aggregations in the Skagerrak have appeared periodically throughout history during the so-called “Bohuslän periods,” in which a large, overwintering stock (9) flooded the Skagerrak such that they could be harvested at scale from the beach (7,8). These periods occurred nearly every century between the 11th and 20th centuries, likely driven by cyclical changes in the North Atlantic Oscillation (NAO) (9). So-called “high” NAO periods cause shifts in current and wind speed resulting in colder temperatures and an influx of North Sea waters (and herring) to the Skagerrak (9). Outside of Bohuslän periods, which typically lasted 20-50 years, NSAS did not appear in the Skagerrak in large enough numbers to approach the coast, and fisheries that sprang up alongside the Bohuslän phenomenon would collapse (8,9). Contemporary NSAS overwinter in the northeastern North Sea on the opposite end of the Norwegian Trench from the Skagerrak (9). This region is also where the Dutch herring industry later started their herring fishing, near Shetland where the largest concentration of herring was to be found prior to- and during their spawning season (4,56). The existence of a NSAS herring in 9th-century Kaupang *prior* to written records thus suggests the Bohuslän herring phenomenon may have occurred at least two centuries earlier than previously known.

### England

The site of Lyminge was an Anglo-Saxon monastery in present-day Kent dating to the 8th century, and is associated with the early medieval revival of marine resource consumption in England, which had largely vanished during the earlier Anglo-Saxon period (57). Our detailed assignment results of herring remains at Lyminge resulted in a mixed signal between Celtic, Downs, and Isle of Man (Figure 3a). Historical evidence suggests herring at Lyminge stem from local populations and were used to feed the religious brethren at the monastery and/or to pay tithes and rent (57). Given the evidence for recent population decline and the known phenomenon of the basin effect in Atlantic herring, the presence of mixed genetic ancestry in Lyminge herring supports our conclusion that herring populations around the UK and Ireland were larger in the past, thus exhibiting a higher degree of genetic connectivity.

Coppergate (10th-11th centuries) is an assemblage from York during the Anglo-Scandinavian Age (42,58). While most of the archaeological specimens could only be assigned to the BINSA metapopulations, several were of high enough quality for detailed assignment, resulting in assignment to mixed origins, including both Celtic/Downs and Isle of Man (Figure 3a). During this period, there is strong evidence of trade between Dublin, the “Southern Isles” (inc. Isle of Man), and York (Jorvik) (59,60). The identification of “Downs” specimens in Coppergate suggest these samples could stem from the East Anglian fishery. East and south coast English herring fisheries are known to have expanded in the 11th century, and historical and archaeological evidence suggests these fish were likely traded across England (6,61). Our genetic results therefore agree with historical sources, but whether the remains from Coppergate were sourced from a single source population, stem from both southern England and Dublin, or changed trade origin over time is a question for future studies with increased power to discriminate between Celtic Sea and Downs populations.

Blue Bridge Lane (BBL) is also an assemblage from York and is dated to the 14th century. These samples come from a single pit full of herring, thus are presumed to represent a single-origin herring shipment (43,58). Detailed genetic assignment of remains from BBL shows that the sample is most related to contemporary Isle of Man (Figure 3a), yet analysis of divergence between temporally-disparate populations suggest it is actually closer to an admixed Celtic/Downs population or the NSAS than contemporary Isle of Man (Figure 3b). Archaeological analysis suggests these remains were not processed like medieval Øresund herring catches, thus they may be from Irish or English fisheries (43,58). At this time, the southern and eastern English fisheries were dominant players in the regional herring market. Fishing towns in Suffolk — one of the counties in East Anglia — dominated the English herring industry in the 10th-14th centuries; Yarmouth, Ipswich, and Dunwich (a now-submerged town) traded fish across England, France, and Germany (11). These towns – Dunwich in particular – are well within current spawning grounds for the Downs population (1,36) and would have targeted coastal spawning aggregations. However, given the high degree of similarity between NSAS and the BBL sample, it is not possible to preclude that this sample stemmed from a local Yorkshire fishery targeting NSAS. Future research on more samples from this site would provide additional clues to specific origin, but our results provide novel genetic evidence (Fig 2) to support the archaeological analysis indicating these samples do not stem from the Øresund herring fishery.

### Netherlands

Huis in den Struys is a site associated with a trading center that started operations in the 13th century in what is now known as Veere, a town in the Zeeland province of the Netherlands (44). Zeeland was known to be a major region practicing distant-water herring fishing for the Dutch Republic by the 15th century (62). The remains are thought to be dated to the 15th century, given that they have been processed for packaging in barrels, which was not locally adopted earlier (44). The Dutch Republic targeted mainly the spawning aggregation on the Fladen Ground off Scotland, but was also known to target the Downs population towards the end of the fishing season from the 14th-17th centuries (4,11). Our detailed assignment of one sample from Huis in den Struys to the Celtic/Downs population provides novel genetic evidence for the Dutch herring industry targeting multiple biological populations. These results highlight the importance of population assignment for modeling historical catch records; the Downs population is clearly genetically distinct from NSAS and has been for over 1000 years. Thus, models using catch records to estimate historical landings must take herring population structure into account to accurately determine historical impacts such as overfishing rather than assuming that all North Sea herring fisheries would have targeted a single, panmictic population.

### Historical implications of temporal genomics analyses

The ancient samples largely exhibited higher individual heterozygosity levels and lower F_ROH_ than contemporary samples, suggesting greater genetic diversity and less inbreeding in the past. The ancient samples further exhibited a wider variation in heterozygosity and in runs of homozygosity. Interestingly, the samples collected in 1979 (NS1, NS2, Downs4(NS)) exhibited levels of heterozygosity more in line with the ancient samples. Given the known impact of low sample sizes and variance in coverage on these analyses (63), these results must be interpreted with caution; larger studies should re-evaluate ROH and heterozygosity in the future. However, we note that we did not find patterns in our results that fit Kardos & Waple’s conclusions that lower coverage is associated with lower F_ROH_ and heterozygosity estimates, suggesting these factors are not driving the variation we see in our results (see Fig 4b,c; Table 1; Supplementary Dataset S1).

Modeling of effective population size over time with GONE revealed strong differentiation between the subpopulations within BINSA, a group that clusters together when conducting structure analysis on the whole genome. GONE is highly sensitive to gene flow and population structure (48), and initial results with incorrect grouping of samples revealed signals of admixture and/or population structure (Figure S9). This further supports the MSA results that there is ongoing population structure between these populations going back through time, particularly between the NSAS and all other BINSA subpopulations. We therefore show that Atlantic herring subpopulations exhibit demographic independence despite known migration between spawning populations (64). This highlights the importance of monitoring these populations as separate management units (28). Further, the demonstrated continuity of population structure throughout the past millennium suggests strong selection for local adaptation.

A recent reevaluation of herring landings across all North Atlantic fisheries since 1500 suggests that catches across all populations hovered around 100,000 tonnes per annum before dramatically growing in the 18th- 19th centuries (10). While fishing pressure across the board was thus relatively constant, pressure on specific stocks/populations shifted dramatically over time. Previous studies on the impact of historical fisheries in the North Sea have suggested several periods of low herring recruitment between 1500 and 2015 were associated with declines in primary productivity (56,65). These fluctuations are not directly reflected in our models of effective population size over time; N_e_ estimates often smooth these relatively short-term census fluctuations (66). We do see, however, a slow decline in N_e_ starting around 1000 CE for the NSAS, which could be due to the combined impact of warmer temperatures (56,67,68) and increasing exploitation (2,4). Intriguingly, this decline slows to seeming stability for NSAS and Isle of Man during the Dutch Herring industry and is not mirrored in the Celtic/Downs admixed population, perhaps due to a larger population size and increased gene flow between the two spawning aggregations.

The Dutch herring period coincided with the onset of the Little Ice Age in the North Sea, which was preceded by the Medieval Warm Period. As previously stated, autumn- and winter-spawning herring inhabit the southern edge of the species range, spawning in warmer waters than any other herring populations. Cool temperatures are also associated with increased growth and fitness for autumn spawning herring (69), suggesting they may have been accompanied by periods of population growth. GONE results do not show population growth, or at least no increase in effective population size, at this time. Historical research indicates exploitation pressure on North Sea herring was below estimated MSY until the late 19th century (4,70), thus it is unlikely to have driven stock collapse. This does not mean, however, there was no impact on SSB or Ne, particularly as we know the Dutch herring industry was targeting spawning aggregations (4). If we presume that a more favorable climate during the Little Ice Age would result in population size increase for the cold-loving NSAS, the Dutch industry may have had a moderating effect on the population. GONE results, which show a stable effective population size in NSAS and Downs herring during the Dutch herring industry, therefore suggest that a stable, favorable climate supports herring stock resiliency. The combination of environmental and anthropogenic factors is also evidenced by further evaluation of the Bohuslän periods. Bohuslän periods occur during cool periods (high NAO index) and are associated with more severe winters (7,71). High NAO indexes have largely disappeared in the late-20th and early 21st centuries due to anthropogenic climate change; the last reported Bohuslän periods lasted from 1877- 1906 (72). The disappearance of Bohuslän periods is likely also affected by overexploitation and poor management drastically reducing the SSB of the source population for this phenomenon (15). Determining the relative impact of each driver is an important question for future studies to explore in order to create lasting sustainable management regimes for this important resource.

The GONE results illustrate declines in N_e_ from five to four million, with one period of slow decline occurring around 1000 CE and a recent period of rapid decline in the last 100 years. While this may appear insignificant – an N_e_ of 4 million and a likely low N_e_/N ratio suggests herring populations are still enormous – it is important to consider this decline in context. First, the most significant portion of this reduction occurred in the past 150 years for all three populations under consideration. This is a near-20% reduction in effective population size, a non-trivial change in the herring genome that could have lasting impacts on its ability to adapt to warming temperatures and maintain crucial population structure and local adaptations. A 2009 study suggested that despite metapopulation recovery in terms of SSB, spatial disparate patterns between the North Sea spawning populations meant the stock did not exhibit full recovery at any point after the 1980s collapse, further finding that the Downs population was contributing more to the North Sea herring population than the NSAS (13). For those populations in our study with samples postdating the recovery period (Celtic/Downs, Isle of Man), we see no recovery in effective population size. Indeed, our discovery of recent divergence between Celtic/Downs population and the Isle of Man population suggests there has been a strong evolutionary impact from the decline through reduced gene flow between these populations. The lack of admixture discovered in these populations in contemporary samples indicates ongoing evolutionary consequences for the Atlantic herring from the overfishing in the 20th century.

Prior studies on the impacts of famous fish stock declines, such as the Grand Banks cod, have found little-to-no effect on genetic diversity (73), although other studies have suggested genetic diversity in marine fish stocks can be impacted without parallel census population decline (74,75). Our own studies on herring indicate that herring are sensitive to overfishing, an observation that is consistent with modeling of the effects of SST change, mass mortality events, and fishing pressure on small pelagic fish species including herring (76–78). Indeed, modeling has shown that small pelagic fish are more vulnerable to environmental changes after exploitation-driven stock collapses (78). The observed reduction in N_e_ is thus likely due to the combined and ongoing stressors of warming temperatures, shifting ocean currents, and exploitation. The dramatic evolutionary consequences highlights the importance of incorporating long-term fishing records and modeling into management advice; allowing a stock to recover somewhat in biomass over the course of a few strong year-classes may not be sufficient to ensure long-term sustainability of these stocks.

### Limitations

This study relies on a relatively small dataset. This is partially due to the poor availability of sequencing data (genome sequences and/or appropriate metadata), with few samples available from prior studies. Despite the small sample size, our reported results are consistent with previous studies and modeling regarding the impact of fisheries and climate change on these herring stocks, and we have successfully recovered finescale population structure with the same loci as reported by Farrell et al. (31) and Bekkevold et al. (28). Demographic modeling with GONE can be conducted with as few as two individuals, although the authors note the exact timing of events may not be entirely accurate (79). However, even with our small sample sizes we recover known historical population declines, suggesting our results are reflecting true processes and not stochastic effects of low sample size. ROH analysis generally requires 4-6 individuals at minimum (80), thus these results should be interpreted with caution. This could explain some of the discrepancies between the ROH results and GONE, such as the lack of evidence of the commercial stock collapse in the 20th century from the ROH results.

The public data for the NSAS was collected in 1979, thus there is a difference of nearly 40 years between sampling for the various BINSA subpopulations in this study. Many of the results presented here seem robust to this difference in sampling time, yet, because this represents up to 10 generations for the NSAS, further studies should endeavor to include more-recently sampled NSAS. Indeed, more recent sampling would better reflect the long-term impact of the stock collapse in the 1960s and the putative ongoing impact of global warming. We have here demonstrated that publicly-available data are incredibly important for molecular ecology work and urge all researchers in this domain to continue to make data available. Increased availability of high-quality data will only strengthen our ability to conduct innovative and robust research on these critical resources. Future studies on these populations that incorporate larger datasets will be well-placed to explore ongoing questions in herring ecology and evolution, such as better characterization of the impacts on population bottlenecks of varying intensity on the herring genome.

### Mixed-stock analysis across a millennium

We were able to assign 8 high-quality (2-23X coverage) ancient genome sequences covering the 8th-15th centuries to local BINSA subpopulations using mixed-stock analysis and found that diagnostic SNPs are robust to long-term change. Atlantic herring exhibit some migration between stocks, although the level of gene flow and interbreeding co-occurring with these trans-stock migrants is unclear (64). We here demonstrate that loci associated with contemporary population structure have changed little over the past millennium, suggesting strong drivers maintaining this structure throughout time. Combined with the result that only autumn spawners were commercially traded, this suggests long-term population stability in the region. These results also show that the creation of diagnostic SNP panels for mixed-stock analysis is a highly cost-effective mode of genetic monitoring, as they can likely be used for decades, if not millennia.

MSA of archaeological remains revealed little differentiation between the Isle of Man and Celtic/Downs populations in the past. Recent studies have shown that these populations exhibit structure, yet have the lowest genetic divergence of all herring stocks thus far assessed using MSA and can be difficult to distinguish based on published markers (31,35). Jost’s D analysis indicates more divergence between contemporary Isle of Man populations compared to the other subpopulations, whereas ancient Isle of Man populations show a strong similarity to Celtic/Downs (Fig 3b). It is unlikely these results are due to low sequence quality, as the ancient specimens were filtered for signatures of contamination, low base quality scores, and strict missingness with the same requirements as the contemporary samples (see Methods). Given the known migration patterns of these three stocks (81), these results suggest possible increased gene flow between the Celtic and Isle of Man populations in the past, with Celtic Sea herring as a mediating population for genetic connectivity between Isle of Man and Downs. Previous research has demonstrated connectivity between Celtic and Downs populations in the 21st century (82), thus, it is likely there was connectivity between these populations in the past when populations were larger.

Increased admixture in the past could have several implications. First, it could indicate that gene flow among these populations is overall too high to fully discriminate between them. It could also indicate that there has been selection pressure or drift on these loci over the last millennium. Celtic Sea and Isle of Man populations were the least-represented samples in the creation of the SNP panels for MSA (28,31). Studies incorporating more contemporary and ancient samples from these populations would greatly strengthen the result by increasing power to differentiate these populations. However, it may also suggest that these loci are capable of discriminating between the subpopulations in contemporary but not ancient samples because there is currently less gene flow than there was in the past. Given known impacts of population decline on herring migration and reproductive patterns (e.g., the basin effect), increased similarity in the past (putative gene flow) is indicative of larger population sizes over the last millennium than seen today in the same region. Indeed, recent evaluation of herring migration patterns in the Celtic Sea concluded that larger population sizes are associated with increased mixing with other stocks and larger migration patterns (81). This suggests that the populations south and west of England should be monitored for overall population health and possible overexploitation. Given the age of the archaeological samples (the most recent with Isle of Man and Downs admixture being from the 15th century), the divergence of these two populations occurred sometime in the past 600 years. If this is associated with population decline, it may be even more recent given the dramatic declines in effective population size in the 20th century recovered with GONE (Fig 4a).

We here demonstrate both the long-term potential of panels developed for mixed-stock analysis of the Atlantic herring and significant changes in herring demography within the last century. Future research efforts with larger, more comprehensive datasets will be able to confirm this pattern, provide more power for discriminating between closely-related subpopulations (e.g., Downs and Celtic Sea), and further explore issues such as exact time of divergence and causal ecosystem dynamics. For example, it has been suggested that Atlantic cod and Atlantic herring do not exist at high biomass at the same time (5) – ancient DNA and genomic modeling on cod and herring simultaneously could reveal shifting stable states in the North Sea between these two economically important species. Better understanding the dynamics at play in the North Sea ecosystem will provide a more solid foundation for developing sustainable management regimes and conservation strategies, such as ensuring ecosystem resilience and the continued survival of fisheries and other species that rely on the Atlantic herring (83).

Our results suggest strong perturbations in herring population health in the 20th century, including likely impacts of climate change and overfishing. Increased admixture between populations in the past is likely due to larger population sizes throughout history compared to modern-day herring stocks, a hypothesis that is supported by the recent population declines recovered through demographic modeling in this study. We find sharp decreases in effective population size in herring populations in the 20th century at a time when overfishing is known to have caused commercial stock collapse in the region. Despite reported increases in census stock size after a brief moratorium on these stocks, effective population size has not recovered for the Atlantic herring stocks here assessed and, indeed, continues to decline. This suggests these populations are not recovering from the 20th- century stock collapses, which could be due to continued overfishing and the effects of warming sea-surface temperatures. We further demonstrate the Atlantic herring exhibit strong population structure that dates back at least 1300 years and can be identified from ancient specimens using the same diagnostic SNP panels that have been developed for contemporary populations. Our detailed assignment from archaeological remains provides novel genetic evidence for historical trade routes and shifting exploitation of various Atlantic herring stocks throughout the last millennium. We further provide novel evidence that extends the known incidence of the so-called “Bohuslän” periods to at least the 9th century, illustrating long-term stability in the region marked by dramatic change in the 20th century. Divergence analysis indicates loci identified for contemporary MSA have changed remarkably little over the past millennium, suggesting strong pressure on these sites to maintain local adaptations and population structure. These results suggest the development of diagnostic SNP panels is a highly cost-effective measure for genetic monitoring of herring stocks, which must be managed with the full knowledge of local adaptations and genetic structure to avoid portfolio effects (29).

## Materials & Methods

### Sampling

Modern data were accessed from publicly available datasets (20,35). The modern dataset comprises 68 individuals from all major herring populations in the Baltic and eastern Atlantic. Archaeological herring material comprising 32 individuals was sourced from 8 sites across Europe spanning the 8th-16th centuries (Figure 2; see Supplementary Dataset S1 for full archaeological context data).

### DNA extraction and sequencing

Archaeological samples were processed following established ancient DNA laboratory protocols to minimize the risk of contamination (84,85). DNA extractions took place at the dedicated ancient DNA laboratory at the University of Oslo. Libraries were prepared with both double-stranded (86) and single-stranded (87) methods. Following AMPure purification and fragment analysis, libraries with sufficient quality were sequenced on an Illumina HiSeq 4000 and/or NovaSeq 6000 at the Norwegian Sequencing Centre. Archaeological samples yielded 0.0001-23X coverage. Coverage for modern sequences ranged from 7-55X. Full metadata for the modern samples is included in Supplementary Dataset S2. All sequences were aligned to the herring reference genome Ch_v2.0.2 (88) using the PALEOMIX (89) v2 pipeline and bwa-mem (90). mapDamage2.0 (91) plots were generated to assess post-mortem deamination patterns and validate the ancient samples (Figure S10).

### Dataset curation

Three datasets were created: modern data, low-coverage ancient and modern data, high coverage ancient and modern data. The modern dataset was used to provide training data and a reference database for assignment tests using BAMscorer. All ancient sequences were used for population assignment tests (see below). Sequences with missingness <30% and coverage >2x were included in a low-coverage dataset in combination with the modern sequences and further processed with angsd (92) (16 ancient and 68 modern samples). High coverage ancient sequences and modern data were combined for further processing using GATK(93) (8 ancient and 68 modern samples). To confirm that all archaeological specimens were Atlantic herring, the mitogenomes were aligned to the reference using GATK for all ancient and modern specimens, filtered for quality, then a maximum-likelihood phylogenetic tree was built with IQ-TREE(94) v1.6.12 (Figure S11). A Pacific herring individual was used as the outgroup (35). All mitogenome sequences were assessed for contamination by examining the rates of heterozygosity with VCFtools (95) after calling loci as pseudo-diploid; as a haploid sequence, non-contaminated individuals should have a heterozygosity rate of 0 using this method. Any individuals with abnormal heterozygosity levels were removed from the dataset.

### SNP Calling

Filtering settings were determined after examining characteristics of the raw VCF files of the final high-coverage and modern datasets, including minor allele frequency, individual depth, locus depth, and locus quality with bcftools (96), VCFtools, and R (97). High coverage datasets were filtered as follows: BCFtools: FS<60.0 && SOR<4 && MQ>30.0 && QD > 2.0 && INFO/DP<415140 –SnpGap 10; VCFtools: --minGQ 15 --minDP 3 -- remove-indels --maf 0.01 --max-missing 0.9; and GATK: –restrict-alleles-to BIALLELIC. The modern dataset contained 4317675 SNPs. For smartPCA and GONE demographic analysis, no maf filtering was applied, as removing minor alleles can artificially reduce signals of population structuring and demographic change (46,98). The non-maf-filtered dataset contained ∼10 million SNPs. An additional dataset was created by removing inversion sites and LD-pruning with PLINKv1.9 (--indep-pairwise 100 10 0.5), resulting in 2846566 SNPs. The high-coverage dataset (including 8 ancient individuals) contained 7 334 073 variants after filtering, with the LD-pruned dataset containing 5673945 variants. For runs of homozygosity, the high-coverage dataset was further reduced to include individuals with <5% missingness, resulting in only two ancient individuals remaining (both from the same context of the 9th century site Lyminge).

### Population structure & MSA (Contemporary)

Population structure was investigated for the non-maf-filtered modern dataset using smartPCA (99). Hudson’s pairwise F_ST_ (100) and kinship coefficients were calculated using popkin (101), both with the full dataset and with a subset of SNPs putatively under selection determined through PCAdapt (102) (Figures S12-13). To further fine-tune population assignment, diagnostic SNPS for mixed-stock analysis (MSA)(28,31) were extracted from the modern and ancient samples identified as belonging to the waters surrounding Britain and Ireland and the North Sea (BINSA). 82 loci in total were called from contemporary samples using angsd (-sites binsa_sites.list - uniqueOnly 1 -remove_bads 1 -only_proper_pairs 1 -trim 0 -C 50 -baq 1 -minMapQ 30 -minQ 20 -noTrans 1 - doCounts 1 -GL 1 -dobcf 1). Individuals with >20% missing loci were removed. Population substructure assessment was conducted in R using DAPC from the adegenet package (37). DAPC was conducted first on the modern data only to confirm the ability of the diagnostic SNPs to assign population as reported in previous studies. The identified subpopulations were used to inform subsequent analysis.

### Population assignment of archaeological specimens

Population assignment tests were conducted on the archaeological samples using BAMscorer v1.6.1(24). Following the methods detailed in Atmore et al. (20), sensitivity analysis was performed for three levels of assignment test based on inversion type and ecological adaptation: salinity adaptation, spawning season, and chr12 inversion haplotype. Spawning season could be determined with as few as 50 000 reads, the chromosome 12 inversion could be assigned with as few as 5 000 reads, and salinity scores could be determined for samples with at least 60 000 reads. All samples were assigned to one of the following populations: British/Irish/North Sea Autumn spawners (BINSA); North East Atlantic Spring spawners (NEAS); Transition zone spring spawners (Transition); Western Baltic Autumn Spawners (WBAS); Central Baltic Autumn Spawners (CBAS); Central Baltic Spring Spawners (CBSS); and Gulf Spring Spawners (GSS) (see Figure 1). All archaeological samples identified as BINSA and with sufficient sequence quality were further investigated with DAPC using the diagnostic SNP panels for mixed-stock analysis. 82 loci in total were called from the ancient high-coverage using angsd (-sites binsa_sites.list -uniqueOnly 1 -remove_bads 1 -only_proper_pairs 1 -trim 0 -C 50 -baq 1 -minMapQ 30 -minQ 20 -noTrans 1 -doCounts 1 -GL 1 -dobcf 1). Individuals with >20% missing loci were removed. To assess the temporal robustness of these diagnostic loci, divergence between the ancient and modern samples were assessed using Jost’s D (45) as a measure of genetic divergence among demes.

### Genetic diversity changes across time

Heterozygosity was measured using angsd on an individual basis for the dataset including ancient samples. To calculate heterozygosity per individual, the individual folded site-frequency spectrum (SFS) was generated using the reference genome as ancestral and standard quality filters (-C 50 -minMapQ 20 -minQ 20 -rmTrans 1). Individual folded SFS were generated with angsd (filters: -uniqueOnly 1 -remove_bads 1 -only_proper_pairs 1 - trim 0 -C 50 -minMapQ 20 -minQ 20 -setMinDepth 5 -setMaxDepth 60 -rmTrans 1) by chromosome. Chromosome-level SFS were then used to calculate individual heterozygosity ranges across the entire genome with realSFS, which takes into account invariant sites.

Runs of homozygosity (ROH) were further explored for all samples with <5% missingness using loci called with angsd and PLINK (see methods). Our previous analysis suggests that ROH estimates are strongly biased by sampling size when overall sampling sizes are small (20), therefore all populations were randomly downsampled to two samples each.Runs of homozygosity (ROH) were assessed using PLINK (103) 1.9 for all samples with <5% missingness with the following commands: --chr-set 26 --double-id --homozyg-snp 50 -- homozyg-kb 90 --homozyg-density 50 --homozyg-gap 1000 --homozyg-window-snp 50 --homozyg-window-het 3 --homozyg-window-missing 10 --homozyg-window-threshold 0.05. To account for the difference in sample size between populations, all populations were sampled to 2 individuals each. Age of coalescent event per ROH was calculated using the following formula: L=100/2g cM (104) (Figure S7).

Treemix (105) was run on the whole-genome dataset with a Pacific herring individual as outlier (missingness set to 0). Treemix showed no divergence between the BINSA subpopulations with the full genome dataset, therefore the diagnostic SNPs from the MSA were extracted from each sample and Treemix was rerun on the smaller dataset (Figures S3, S5). Treemix was bootstrapped through 10 iterations per migration edge (0–5) and then an optimal migration edge was selected using OptM (38) (Figure S5). The optimal migration edge was then used as a setting for bootstrapping and visualization with BITE (106).

### Demographic modeling

The non-maf-filtered modern dataset was used to estimate effective population size throughout time with GONE (46). The BINSA population was assessed both as a homogenous population and by geographic sampling location to determine the possibility of cryptic population substructure and to control for the difference in sampling time between the oldest and newest specimens (∼50 years = up to 15 generations). GONE results were used to confirm mixed-stock analysis with DAPC by assessing the presence or absence of artifacts known to be produced by admixture (48) (Figures S8-9). Known inversions (35) were removed from the dataset prior to running GONE. GONE internally bootstraps Ne estimation through 40 iterations for each generation using a random subsampling of 50 000 SNPs from each chromosome per iteration. The program provides a geometric mean population size per gen. To assess reliability of the program given known sensitivity to low sample sizes (46), each population from the modern dataset was run through GONE 30 times, with the median population size estimate per gen taken as the consensus effective population size at a given time depth. Default parameters were used other than including the recombination rate for the Atlantic herring (2.54 cM/Mb) (35,88). Time depth was estimated using 3-5 years as a minimum generation time for Atlantic herring based on average generation time per subpopulation computed from Feng et al. (107).

## Author Contributions

L.M.A wrote the paper with contributions from B.S. L.M.A., B.S., J.H.B. conceived of the study. L.M.A. designed the study and conducted analysis with input from B.S. Lab work was conducted by L.M.A. and A.B. Archaeological samples and contextual information were provided by I.J., S.H., A.S., J.G., R.B., K.D., L.Q., D.C.O., and A.K.H. All authors agreed on the final text for this paper.

## Acknowledgements

We are very grateful to the York Archaeological Trust and the Cultural Heritage Agency of the Netherlands for allowing us to conduct destructive analysis on archaeological herring remains in their collections. We also extend our gratitude to Cecily Spall from Field Archaeology Specialists Heritage, and Joran Smale from the Provincial Archaeological Depot in Flevoland for access to samples from Blue Bridge Lane and the Biddinghuizen shipwreck. Finally, we thank Professor Gabor Thomas for providing access to the archaeological samples from Lyminge. This project has received funding from the European Union’s Horizon 2020 research and innovation program under Marie Skłodowska-Curie Grant Agreement No. 813383 (Seachanges) and from the European Research Council under the European Union’s Horizon 2020 research and innovation program (4-OCEANS, Grant Agreement No. 951649).

