## Supplement for "The Once and Future Fish: 1300 years of Atlantic herring population structure and demography revealed through ancient DNA and mixed-stock analysis"

**This Supplement Includes:**

Supplementary figures S1-S13

**Supplementary Figures**

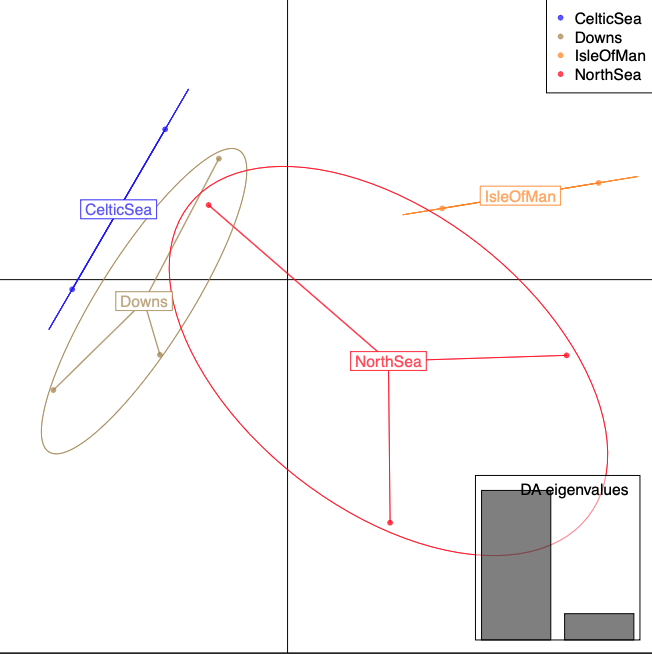

**Figure S1** – **PCA produced with DAPC** suggesting one individual from the North Sea population clusters with the Downs population using SNPs designed to discriminate between Downs and North Sea herring (24).

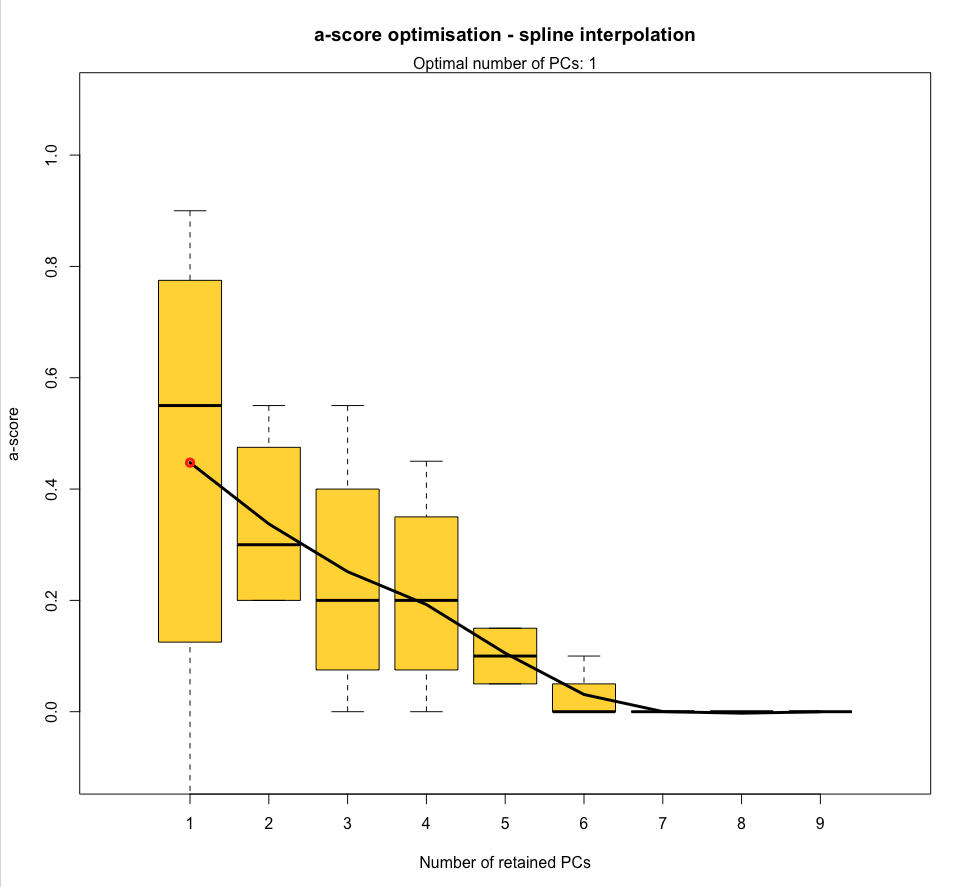

**Figure S2 – Spline analysis for optimal number of PCs to retain for DAPC analysis.** Results suggest using one PC for discriminant analysis.

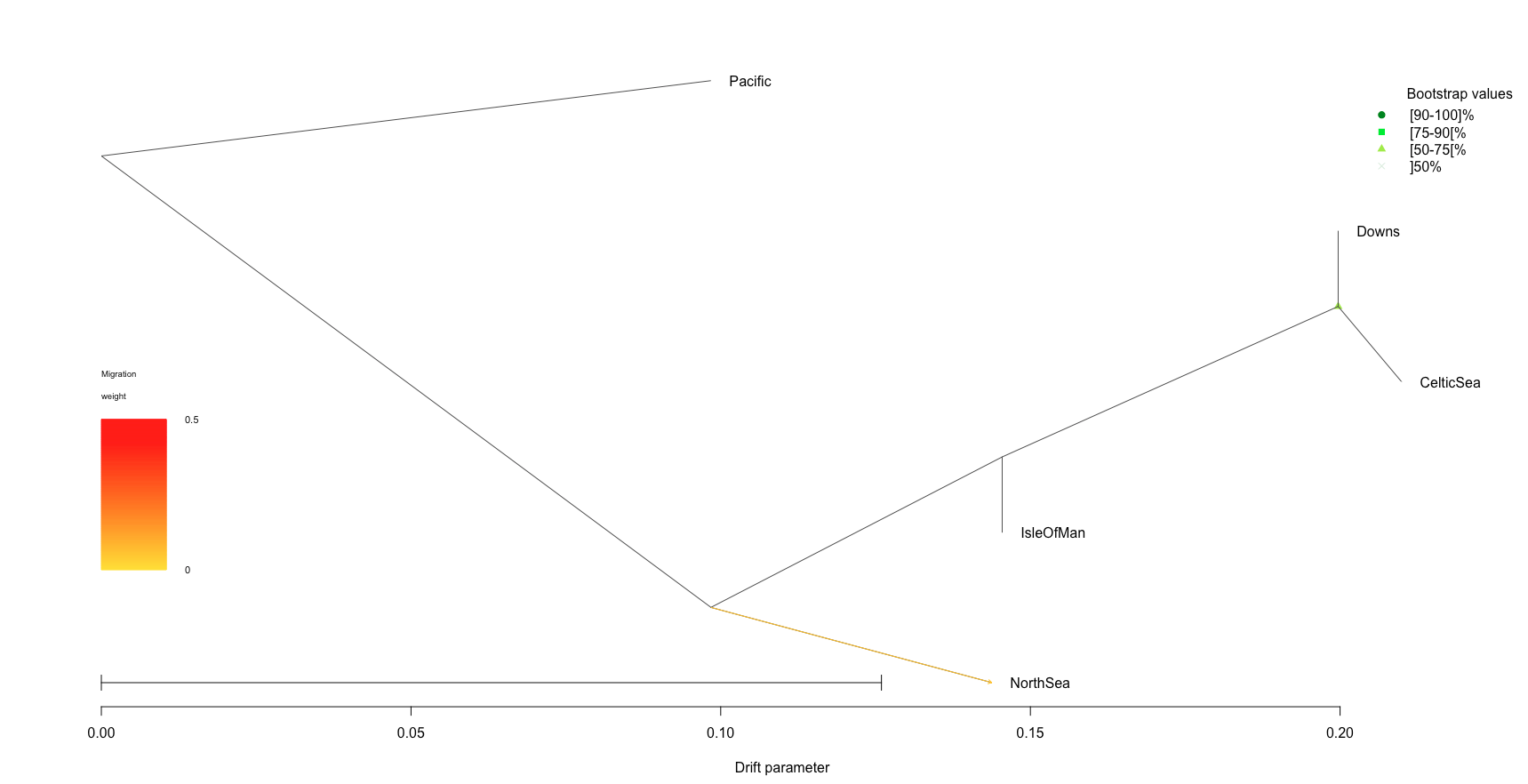

**Figure S3 – Treemix with 1 migration edge for MSA diagnostic SNPs** on contemporary samples suggests separation between North Sean and Downs/Celtic/Irish seas, with Downs and Celtic Sea being most closely-related. The single migration edge appears from the common ancestor of the western BINSA populations to the North Sea. All nodes have 100% bootstrap support.

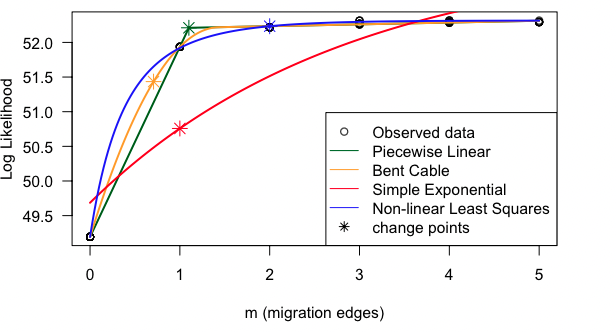

**Figure S4 – OptM results** suggesting one migration edge is appropriate for the MSA diagnostic SNP data

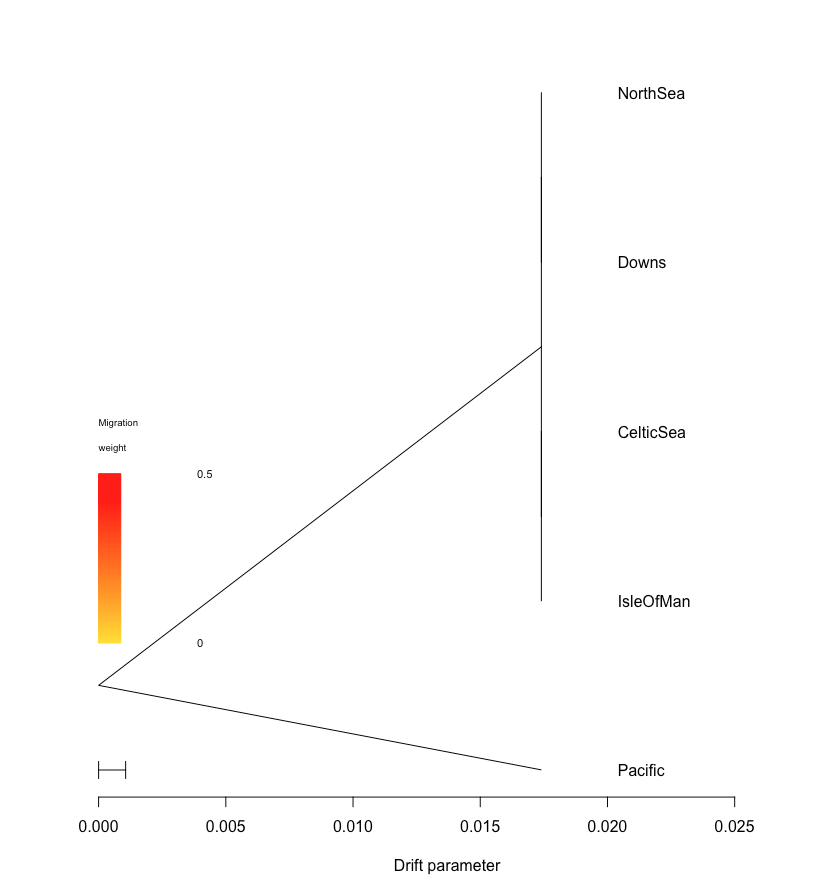

**Figure S5 – Treemix results for LD-pruned whole-genome dataset** on contemporary samples reveals no population structure between the BINSA populations.

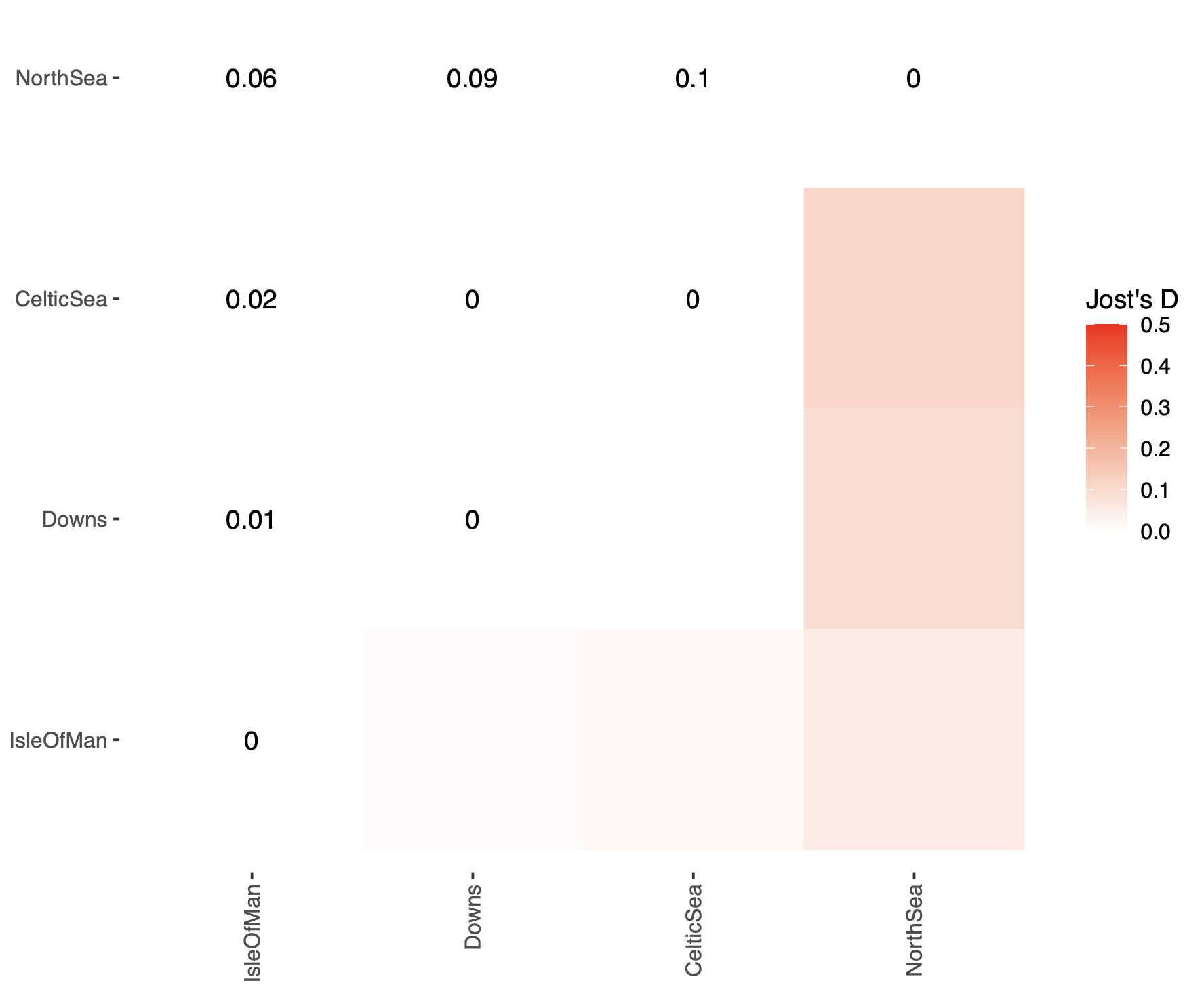
**Figure S6 – Jost’s D analysis of divergence between modern BINSA subpopulations** showing no differentiation between Downs and CelticSea and some variation between this population and Isle of Man. The majority of differentiation occurs between the North Sea and all other populations.

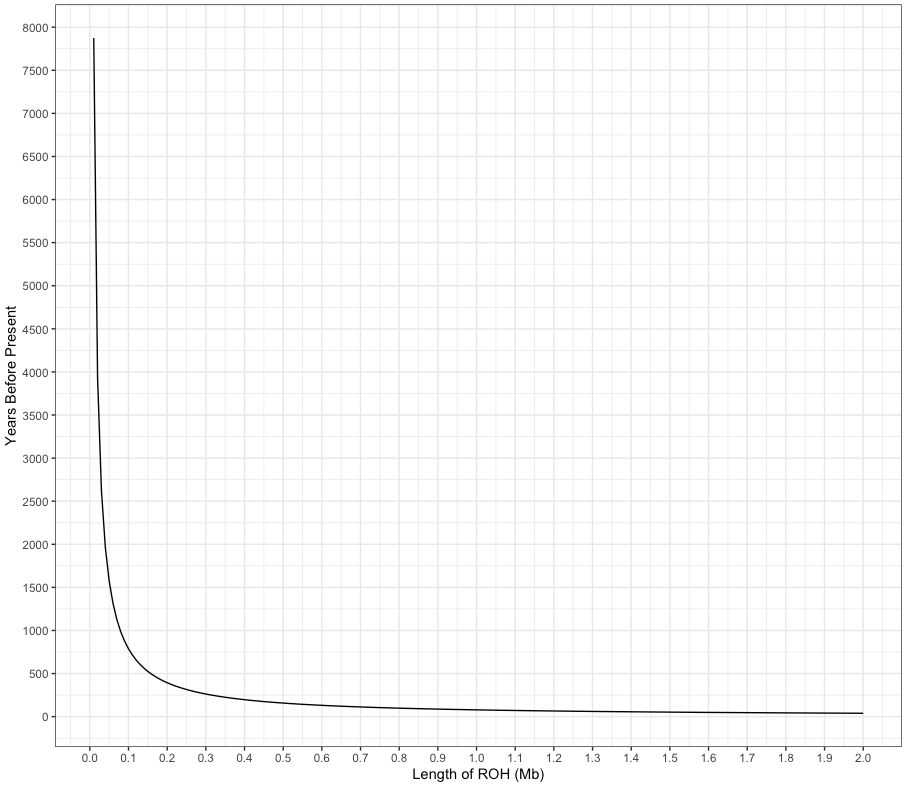

**Figure S7 – Time to coalescence for length of Run of Homozygosity.** Estimated using the formula 100/2g cM/Mb=L and herring recombination rate of 2.54 cM/Mb (Pettersson et al., 2019) and generation time of 4 years.

**Figure S8 – GONE on BINSA metapopulation combined** shows many different population trajectories for each iteration, indicating possible population structure confounding the signal.
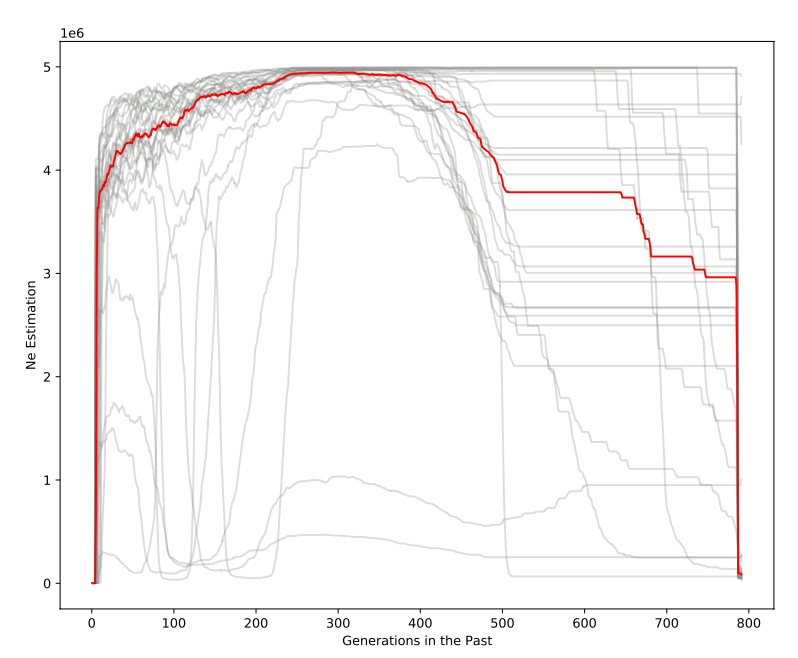

a)	North Sea	b) Downs
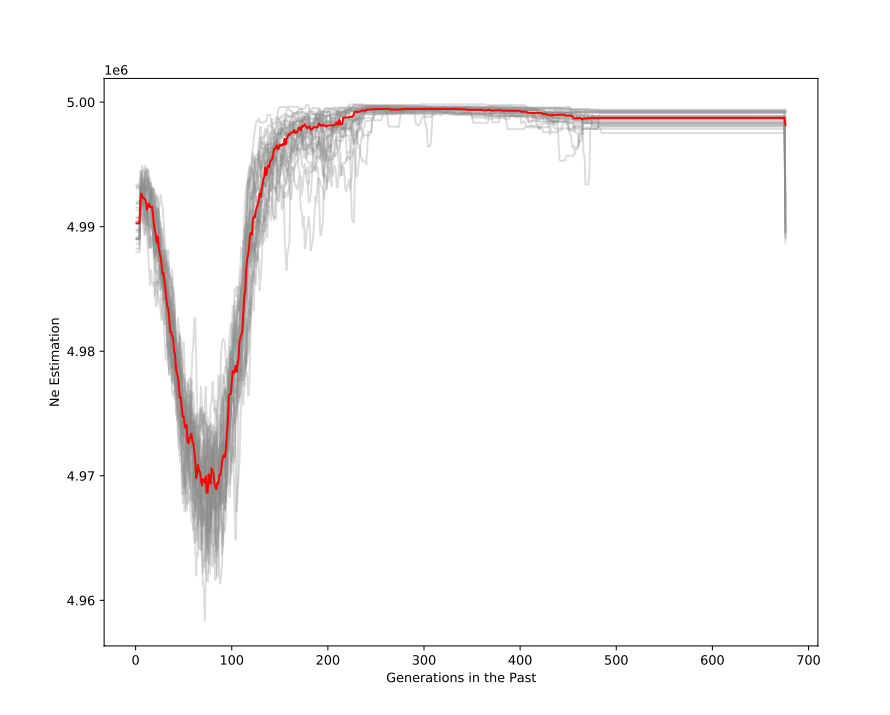

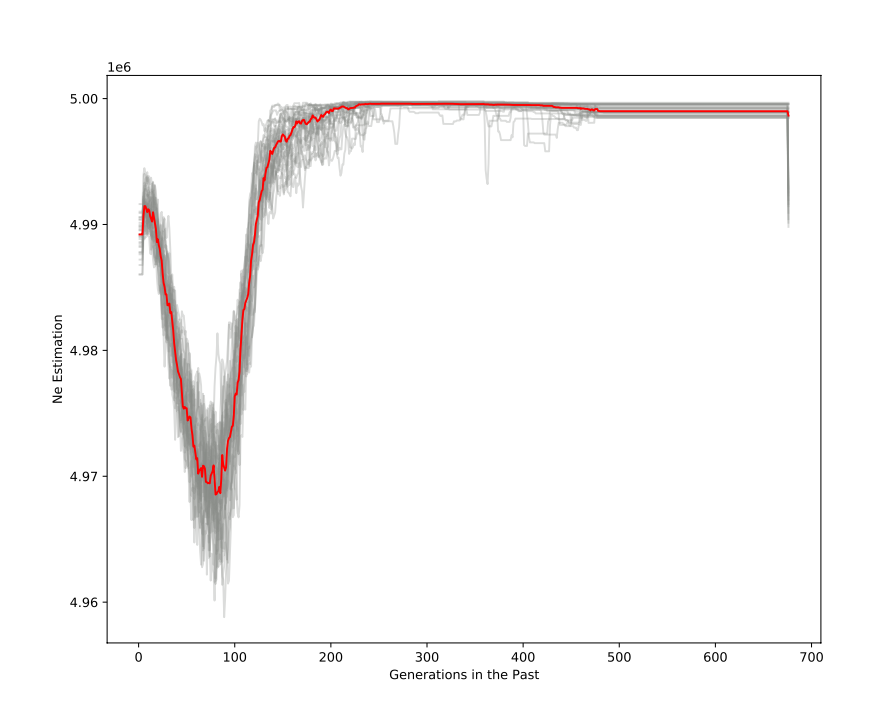

c) Celtic Sea	d) Isle Of Man (Irish Sea)

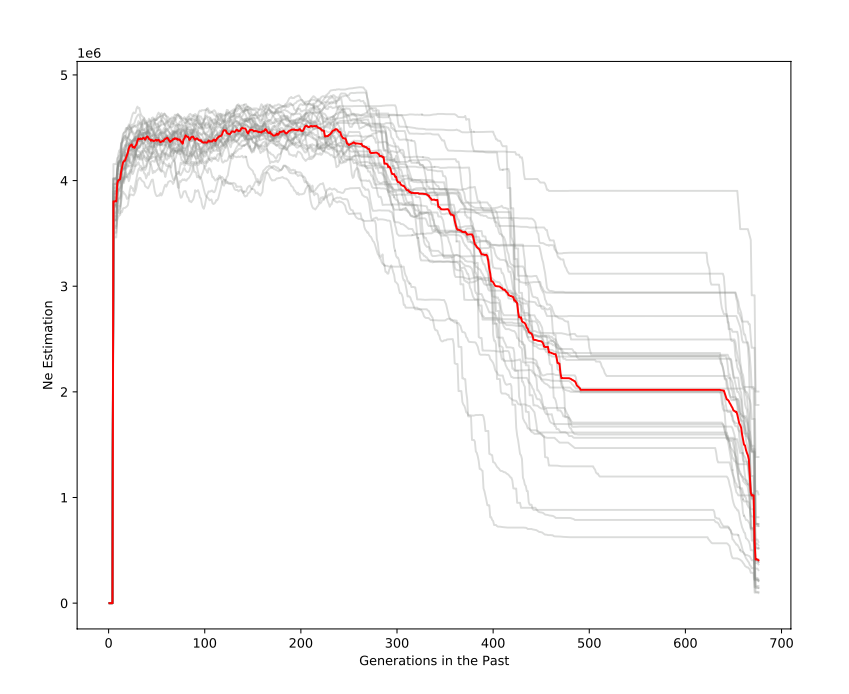

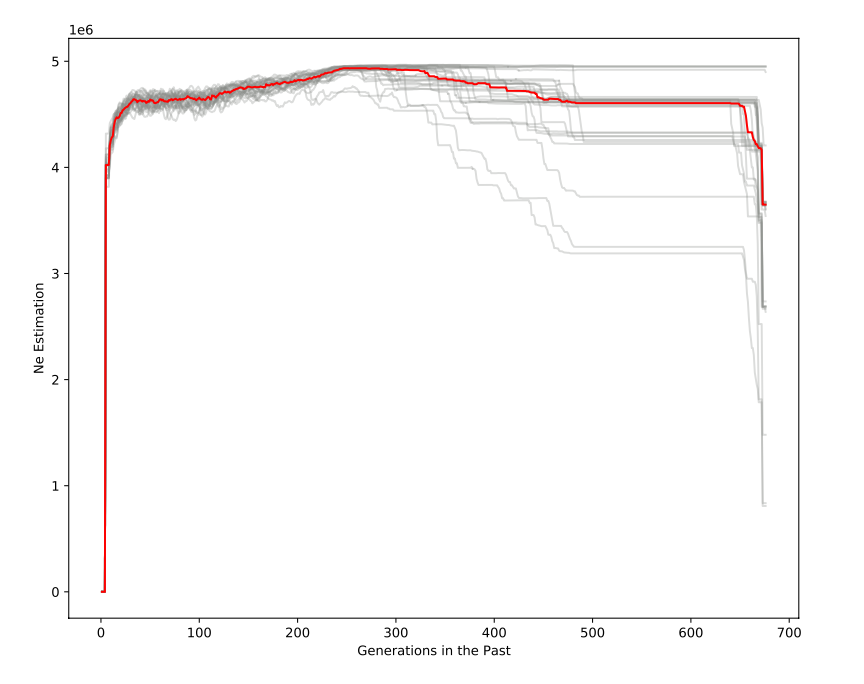

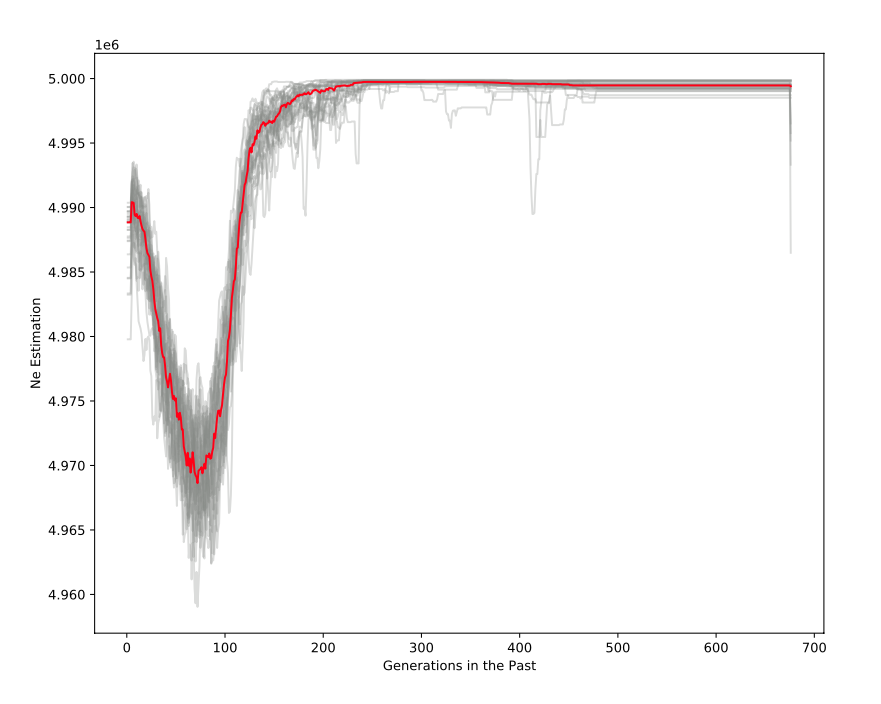

e) Downs with NorthSea13

**Figure S9 – GONE run on each BINSA subpopulation split by sampling location** (according to downloaded metadata) suggesting undetected admixture in Downs and North Sea populations. Combined Downs and NorthSea13 still show signs of admixture, likely because NorthSea13 was sampled in 1979 as opposed to 2016 for Downs.

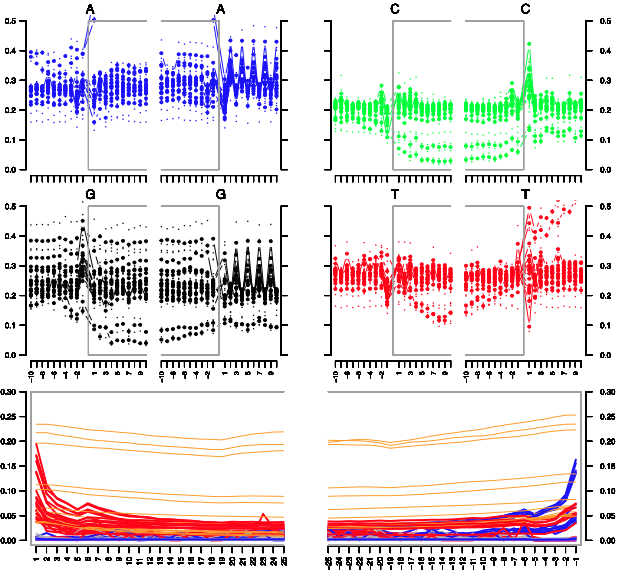

**Figure S10 – mapDamage plots for ancient individuals.** Ancient samples exhibited classic signs of postmortem damage, which validates their interpretation as ancient samples. Those samples exhibiting contamination were removed from analysis.

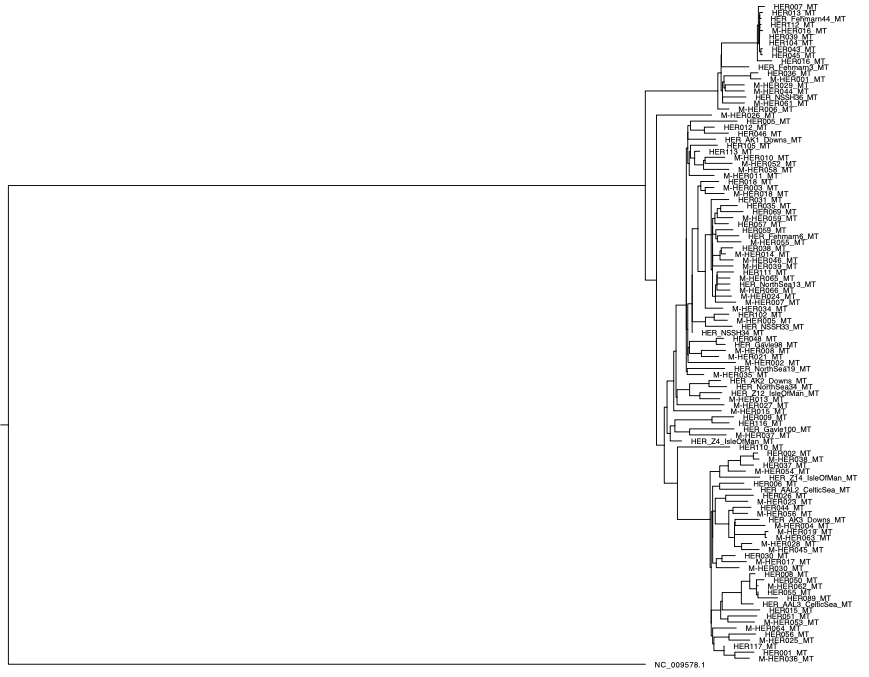

**Figure S11 – Maximum Likelihood Tree for Herring Mitogenome.** Constructed using IQ-Tree and a Pacific herring (*Clupea pallasii*) as the outgroup, this tree confirms the ancient samples fall within the diversity of Atlantic herring. No mitogenome structure corresponding to geography is exhibited.

**a)**
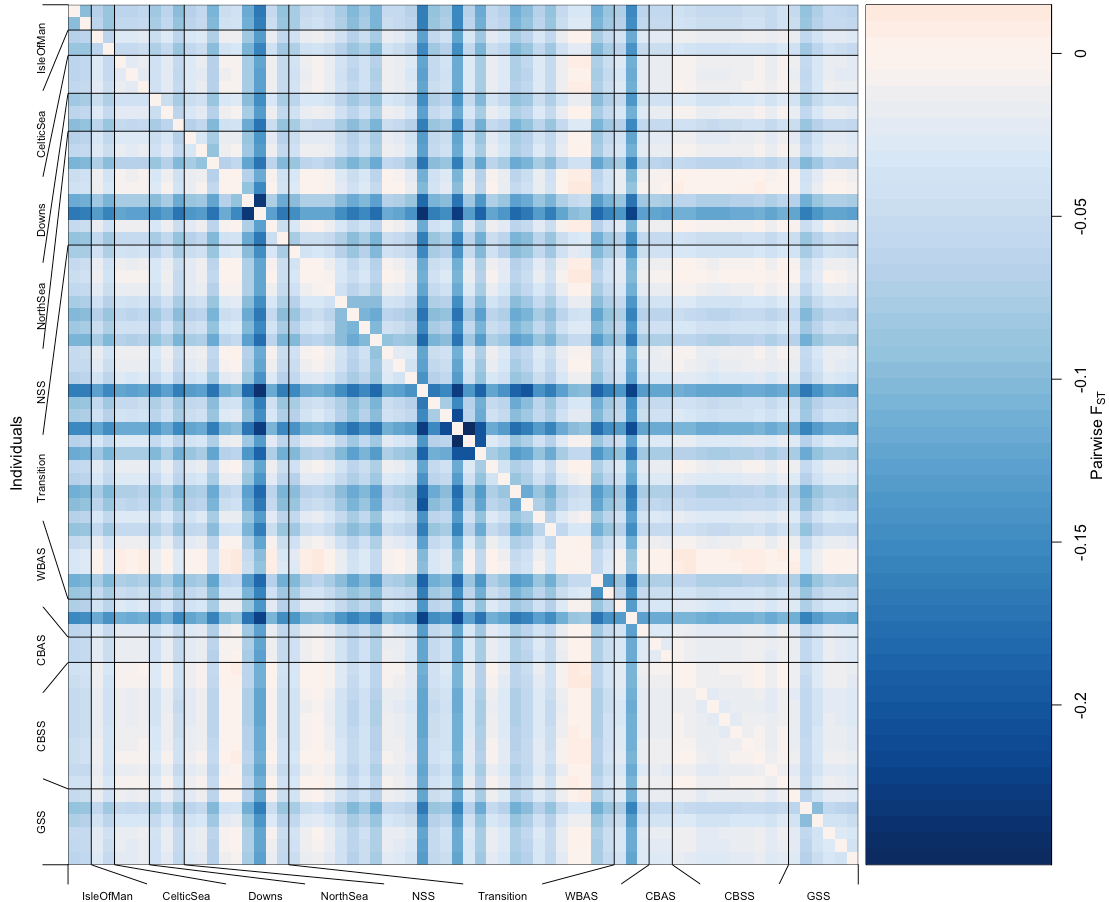

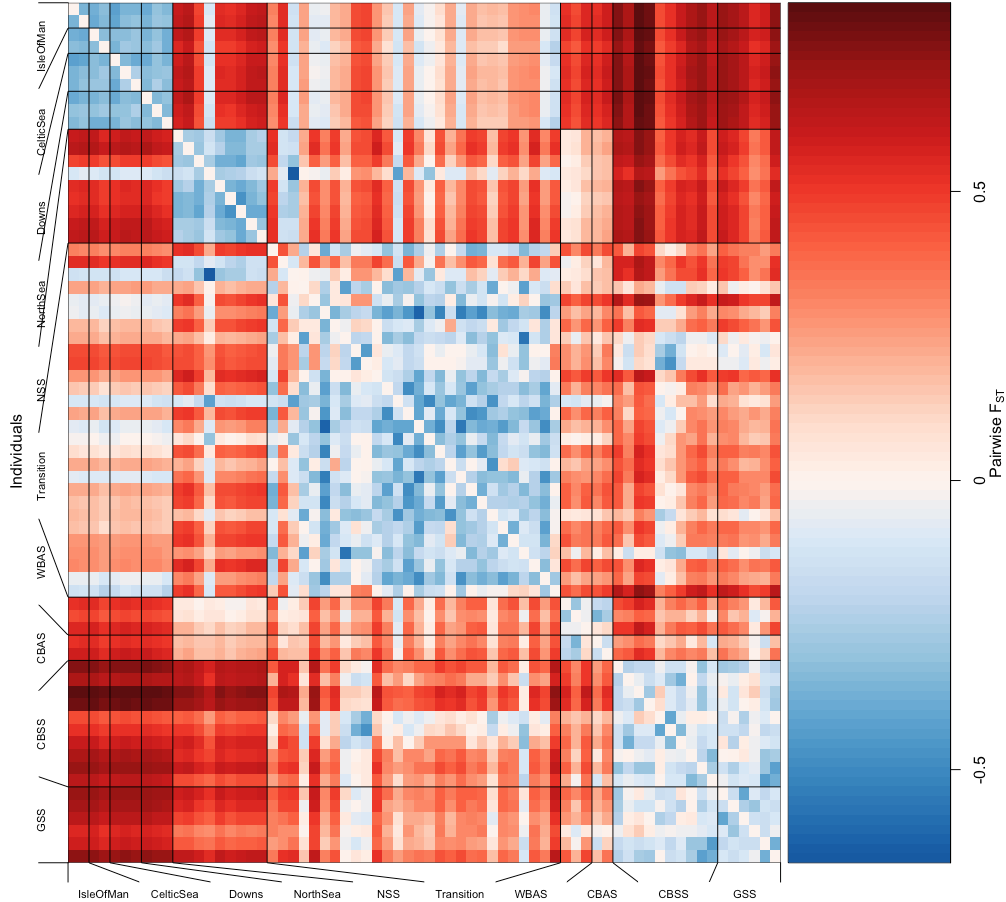

**b)**

**Figure S12 – Pairwise Fst with Popkin.** a) Hudson’s pairwise Fst calculated with neutral SNPs (~4 million, maf-filtered and pruned for LD) from 68 contemporary herring samples across eastern Atlantic and Baltic. Individual pairwise Fst estimates show zero or negative Fst values, indicating no population structure across all Atlantic herring populations in this study; b) Hudson’s pairwise Fst with SNPs identified as outliers with PCAdapt. Negative or zero Fst values are here restricted to within-group variation, while between group variation is delineated based on spawning season and adaptation to salinity reflecting known population structure

**a)**
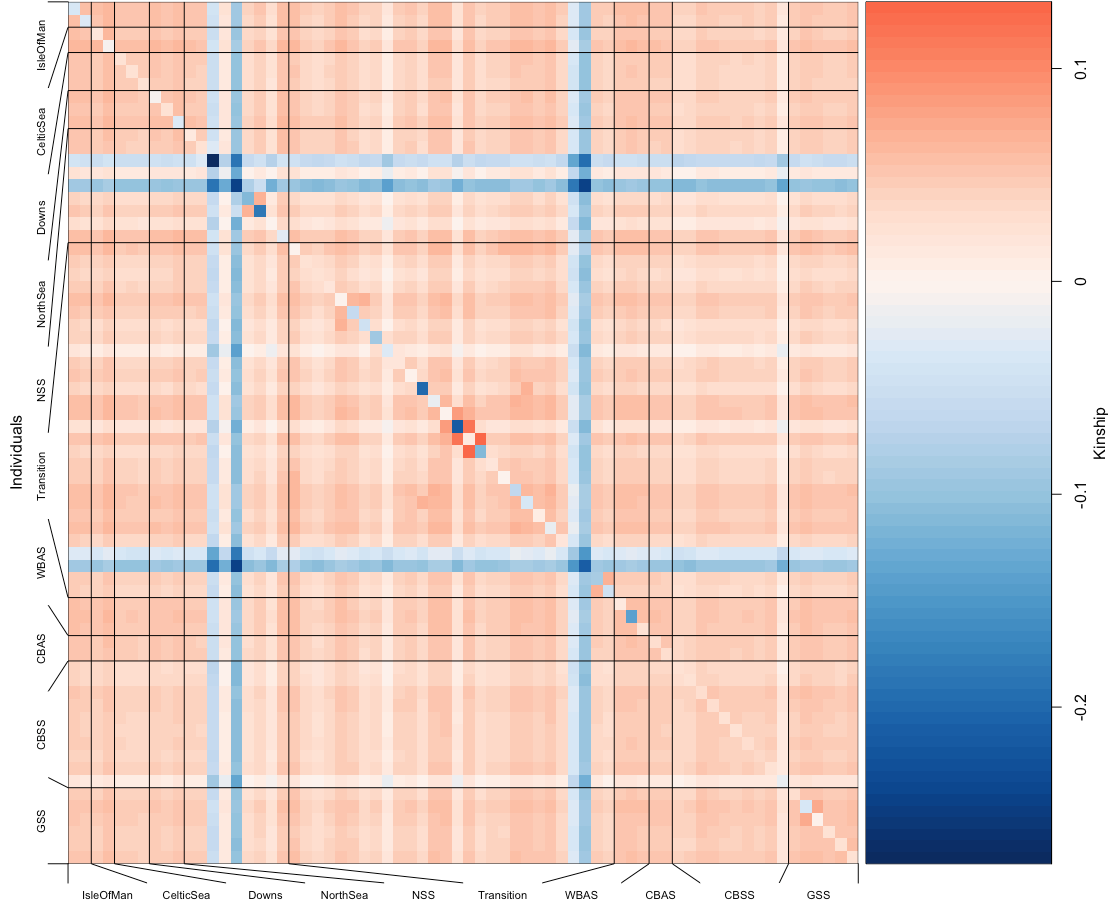

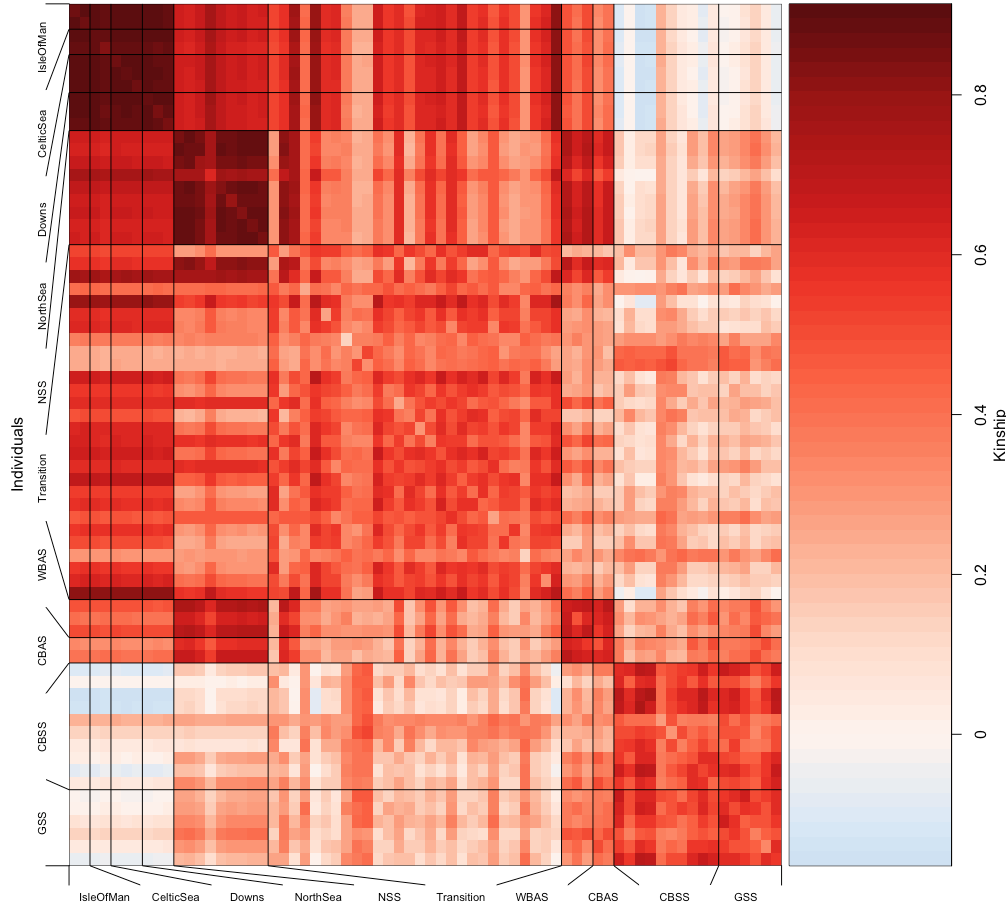

**b)**

**Figure S13 – Pairwise individual kinship matrix.** The kinship matrix was calculated using popkin on 68 contemporary herring samples from across the eastern Atlantic and the Baltic. a) ~4 million neutral SNPs (MAF-filtered, LD-pruned) fail to recover any population structure across these metapopulations; b) SNPs identified as outliers by PCAdapt show increased kinship within known metapopulations delineated by salinity adaptation and spawning season.
